# Surface-based Single-subject Morphological Brain Networks: Effects of Morphological Index, Brain Parcellation and Similarity Measure, Sample Size-varying Stability and Test-retest Reliability

**DOI:** 10.1101/2021.01.25.428021

**Authors:** Yinzhi Li, Ningkai Wang, Hao Wang, Yating Lv, Qihong Zou, Jinhui Wang

## Abstract

Morphological brain networks, in particular those at the individual level, have become an important approach for studying the human brain connectome; however, relevant methodology is far from being well-established in their formation, description and reproducibility. Here, we extended our previous study by constructing and characterizing single-subject morphological similarity networks from brain volume to surface space and systematically evaluated their reproducibility with respect to effects of different choices of morphological index, brain parcellation atlas and similarity measure, sample size-varying stability and test-retest reliability. Using the Human Connectome Project dataset, we found that surface-based single-subject morphological similarity networks shared common small-world organization, high parallel efficiency, modular architecture and bilaterally distributed hubs regardless of different analytical strategies. Nevertheless, quantitative values of all interregional similarities, global network measures and nodal centralities were significantly affected by choices of morphological index, brain parcellation atlas and similarity measure. Moreover, the morphological similarity networks varied along with the number of participants and approached stability until the sample size exceeded ∼70. Using an independent test-retest dataset, we found fair to good, even excellent, reliability for most interregional similarities and network measures, which were also modulated by different analytical strategies, in particular choices of morphological index. Specifically, fractal dimension and sulcal depth outperformed gyrification index and cortical thickness, higher-resolution atlases outperformed lower-resolution atlases, and Jensen-Shannon divergence-based similarity outperformed Kullback-Leibler divergence-based similarity. Altogether, our findings propose surface-based single-subject morphological similarity networks as a reliable method to characterize the human brain connectome and provide methodological recommendations and guidance for future research.

## Introduction

Morphological brain networks depict patterns of interregional relations in regional brain morphology on the basis of structural magnetic resonance imaging. Historically, morphological brain networks are mainly derived via population-based morphological covariance network methods by estimating interregional covariance across a cohort of participants in a certain morphological index, such as gray matter volume, cortical thickness and surface area (Bassett et al., 2008; He et al., 2007; Sanabria-Diaz et al., 2010). To date, the population-based morphological covariance network methods have been widely used as an important tool to study the human brain, including but not limited to parsing of organizational principles of healthy brains, characterization of trajectories during development and aging and identification of abnormalities in various brain diseases (see Alexander-Bloch et al., 2013a; Evans, 2013 for two excellent reviews).

However, the population-based morphological covariance networks suffer from several noticeable issues, such as neglect of interindividual variability, requirement of a large sample size and introduction of complicated models for subsequent statistical inference. All these issues limit the universal application of morphological brain networks, in particular in uncovering their neurobiological significance and clinical diagnostic and prognostic value. Recently, the advent of individual-level morphological similarity networks has overcome, to a great extent, these issues and has thus attracted considerable attention (Jiang et al., 2017; Kong et al., 2015; Li et al., 2017; Seidlitz et al., 2018; Tijms et al., 2012; Wang et al., 2016; Yu et al., 2018). With these individual-level methods, several studies show that morphological similarity networks can capture known cortical cytoarchitecture and related gene expression (Seidlitz et al., 2018), account for interindividual differences in cognition (Li and Kong, 2017; Seidlitz et al., 2018; Tijms et al., 2014), and distinguish patients with schizophrenia from healthy controls (Zhao et al., 2020) and predict clinical progression of patients with Alzheimer’s disease (Tijms et al., 2018). These findings provide strong evidence that individual-level morphological similarity networks are biologically meaningful and are of great value in helping clinical diagnosis and prognosis.

Technically, individual-level morphological similarity networks can be divided into two subcategories. The first estimates interregional morphological similarity by computing Pearson correlation coefficients in regional mean signals across different morphological indices (Li et al., 2017; Seidlitz et al., 2018). This type of method, however, not only neglects intraregional morphological distributions but also may lead to unstable similarity estimation due to a limited number of morphological indices available (< 10 in previous studies), which are treated as samples for the correlation analysis. In contrast, the second subcategory estimates interregional morphological similarity at a more refined level by taking intraregional morphological distributions into account from different perspectives (Jiang et al., 2017; Kong et al., 2015; Tijms et al., 2012; Yu et al., 2018). In this regard, we previously constructed single-subject morphological similarity networks by estimating interregional morphological similarity in the distribution of regional gray matter volume in terms of the Kullback-Leibler divergence (*KLD*; Wang et al., 2016). We found that this method can reveal structured organization of morphological similarity networks with high test-retest (TRT) reliability (Wang et al., 2016). It should be noted that the Pearson correlation-based method and the *KLD*-based method are essentially different: the former estimates interregional morphological similarity in intraregional mean of multiple morphological indices, while the latter estimates interregional morphological similarity based on intraregional distribution of a single morphological index. The Pearson correlation method cannot be used to estimate interregional morphological similarity based on a single morphological index. This is not only because there are different numbers of vertices but also because there is no one-to-one correspondence of the vertices between two regions. Thus, these two types of methods cannot be compared directly.

In the present contribution, we extended our previous study in several aspects. First, we constructed single-subject morphological similarity networks in cerebral cortical surface rather than in volume space. The greatest benefit from this switch is that spherical registration of cortical surface meshes increases the accuracy of brain registration and thus allows more precise localization (Desai et al., 2005). In addition, inflation of cortical surface meshes promotes better visualization of buried sulci by raising them to the surface. Second, there are more forms of morphological indices available for surface mesh-based analysis, with each capturing different aspects of cerebral morphology. Thus, in contrast to a single morphological index (i.e., gray matter volume), used in our previous study, this work examined common and specific degrees of organization among different types of morphological similarity networks derived from different morphological indices. These analyses are of great significance for comprehensive understanding of morphological similarity network architecture. Third, in addition to evaluating different brain parcellation atlases as in our previous study, this work examined effects of different similarity measures on morphological similarity networks. Despite accumulating evidence for significant impacts of different similarity measures on structural and functional brain networks (Liang et al., 2012; Sarwar et al., 2019; Zalesky et al., 2010), the extent to which morphological similarity networks depends on choices of similarity measure is largely unknown. Finally, this work evaluated stability with respect to different sample sizes and TRT reliability of morphological similarity networks.

We hypothesize that quantitative descriptions of surface-based single-subject morphological similarity networks are dependent on choices of morphological index, brain parcellation atlas and similarity measure, are robust to variation in sample size and have high TRT reliability.

## Materials and Methods

### General analytical pipeline

In this study, using a large-scale dataset, we first derived 4 surface-based, vertexwise morphological brain maps (fractal dimension, FD, gyrification index, GI, sulcal depth, SD, and cortical thickness, CT) for each participant, based on which single-subject morphological similarity networks were constructed using two surface atlases for brain parcellation (a2009s atlas and a2005s atlas) and two similarity measures for interregional similarity estimation (*KLD*-based similarity, *KLDs*, and Jensen-Shannon divergence-based similarity, *JSDs*). Therefore, we obtained 16 morphological similarity networks in total for each participant (4 morphological indices × 2 parcellation atlases × 2 similarity measures). Then, we examined effects of the different analytical strategies as well as sample sizes on interregional similarity, global network organization and nodal centrality of the resultant morphological similarity networks. Finally, we utilized an independent, TRT dataset to explore reliability of surface-based single-subject morphological similarity networks under different analytical strategies, aiming to provide methodological recommendations for future studies. Figure 1 presents a flowchart of the overall pipeline of data processing.

**Figure 1.**
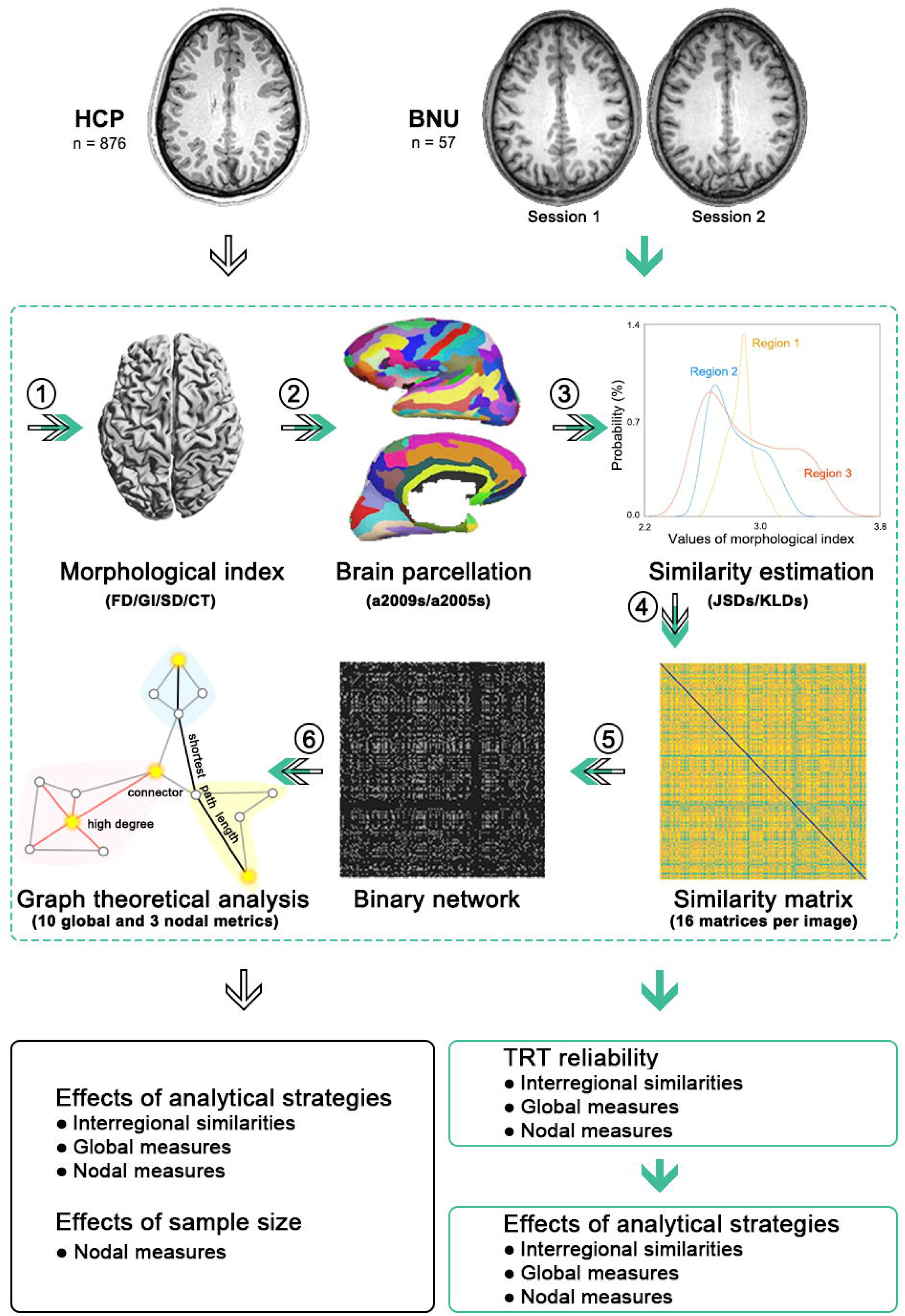
Flow chart of imaging processing, network construction, topological characterization and statistical analysis in this study. ➀ For each structural image from the HCP and BNU datasets, four vertexwise morphological maps (FD, GI, SD and CT) were first extracted. ➁ Each morphological map was then divided into different numbers of regions according to two brain atlases (a2009s and a2005s). ➂ Subsequently, the probability distribution function was estimated for each region in terms of signal distribution of each morphological index and was used to estimate interregional similarity with *JSDs* and *KLDs*, respectively. ➃ This formed a total of 16 similarity matrices for each image (4 morphological indices × 2 brain parcellation atlases× 2 similarity measures). ➄ Before topological characterization of the resultant similarity matrices, a sparsity-based procedure was further used to threshold each of them into a series of binary networks. ➅ Finally, 10 global and 3 nodal graph theory-based network metrics were calculated to characterize topological organization of each binary network. FD, fractal dimension; GI, gyrification index; SD, sulcal depth; CT, cortical thickness; *JSDs*, Jensen-Shannon divergence-based similarity; *KLDs*, Kullback-Leibler divergence-based similarity; TRT, test-retest.

### Participants

Two publicly available datasets were used in this study: the Human Connectome Project (HCP) S900 dataset (www.humanconnectome.org) (Van Essen et al., 2013) and the Beijing Normal University (BNU) TRT dataset (http://fcon_1000.projects.nitrc.org/indi/CoRR/html/bnu_1.html). The former was used to characterize the topological organization of morphological similarity networks under different analytical strategies and effects of different sample sizes, and the latter was used to evaluate TRT reliabilities of morphological similarity networks.

#### HCP S900 dataset

The HCP S900 dataset includes a total of 897 healthy adult participants who completed structural MRI scans. Twenty-one participants were excluded according to our quality control procedures (see below), resulting in 876 participants in the final analyses (male/female: 386/490; main age: 22-35 years).

#### BNU TRT dataset

The BNU TRT dataset contains a total of 57 healthy young participants (male/female: 30/27; age: 19-30 years) who each completed two MRI scan sessions within an interval of approximately 6-weeks (40.94 ± 4.51 days). All participants were right-handed and had no history of neurological or psychiatric disorders.

### MRI data acquisition

#### HCP S900 dataset

T1w images from the HCP S900 dataset were obtained using a customized 3T Siemens Magnetom Connectome scanner with a 32-channel head coil. Main imaging parameters were: repetition time (TR) = 2400 ms, echo time (TE) = 2.14 ms, inversion time (TI) = 1000 ms, flip angle (FA) = 8°, field of view (FOV) = 224 × 224 mm^2^, matrix = 320 × 320, thickness = 0.7 mm with no gap, and 256 sagittal slices.

#### BNU TRT dataset

T1w images from the BNU TRT dataset were obtained using a 3T Siemens Tim Trio scanner with a 12-channel head coil. Main imaging parameters were: TR = 2530 ms, TE = 3.39 ms, TI = 1100 ms, FA = 7°, FOV = 256 × 256 mm^2^, matrix = 256 ×192, slice thickness = 1.33 mm; interslice gap = 0.65 mm, and 144 sagittal slices.

### Quality control procedures

The HCP S900 dataset underwent multiple levels of quality control, ranging from real-time oversight during acquisition to post-acquisition manual and automated image review. Each of these procedures is specified in a formal standard operating procedure and integrated into the internal database system (Marcus et al., 2013). For the BNU TRT dataset, the quality control procedures included visual inspection of severe motion artefacts or any other apparent artefacts, followed by calculation of a series of quality evaluation metrics, such as signal-to-noise ratio, foreground to background energy ratio, ghost to signal ratio and artifact detection (Lin et al., 2015). In addition to the dataset specific quality control procedures, in this study we further visually checked the results of image segmentation via the modules “Slice Display” and “Surface Data Homogeneity” in the CAT12 toolbox. Twenty-one participants were excluded due to failed segmentation or poor image quality for the HCP S900 dataset.

### MRI data preprocessing

All structural images underwent standard processes using Computation Anatomy Toolbox 12 (CAT12, version r1113, http://www.neuro.uni-jena.de/cat/), based on Statistical Parametric Mapping 12 (SPM12, version 6685, https://www.fil.ion.ucl.ac.uk/spm/software/spm12). The CAT12 offers a volume-based approach for estimating cerebral surface morphology without extensive reconstruction of cortical surface and thus is timesaving. Specifically, the CAT12 contains a processing pipeline for computing four morphological indices, including FD, GI, SD and CT. All image preprocessing and morphological parameter computation described below were conducted in subject native space. After obtaining individual morphological maps of FD, GI, SD and CT, they were finally resampled into the common fsaverage template and smoothed using a Gaussian kernel. Specifically, individual CT maps were smoothed using a Gaussian kernel with 15-mm full width at half maximum, while individual FD, GI and SD maps were smoothed using a Gaussian kernel with 25-mm full width at half maximum. Based on the recommendations of the CAT12 manual, the usage of larger filter sizes for FD, GI and SD is due to the underlying nature of these folding measures that reflect contributions from both sulci and gyri. Therefore, the filter size should exceed the distance between a gyral crown and a sulcal fundus.

#### CT calculation

CT was estimated using a fast and reliable projection-based thickness (PBT) method, which requires no extensive reconstruction of the cortical surface. First, following tissue segmentation, a white matter (WM) distance map was derived by estimating the distance from the inner gray matter (GM) boundary for each GM voxel (Dahnke et al., 2013). Values at the outer GM boundary in the WM distance map (i.e., GM thickness) were then projected back to the inner GM boundary to generate a GM thickness map. Subsequently, a central surface was created at the 50% level of the percentage position between the WM distance and GM thickness maps. For the resultant central surface, a topology correction based on spherical harmonics was used to account for topological defects (Yotter et al., 2011a). Furthermore, the central surface was reparameterized into a common coordinate system via spherical mapping (Yotter et al., 2011c), and spherical registration adopted the volume-based diffeomorphic DARTEL algorithm (Ashburner, 2007) to the surface.

#### FD calculation

FD reflects cortical surface folding complexity. It was estimated based on spherical harmonic reconstructions and calculated as the slope of a logarithmic plot of surface area versus the maximum l-value, where the maximum l-value is a measure of the bandwidth of frequencies used to reconstruct the surface shape (Yotter et al., 2011b).

#### GI calculation

GI is an indicator of cortical folding. Based on the spherical harmonic reconstructions, GI was calculated as absolute mean curvature (Luders et al., 2006). Mean curvature is an extrinsic surface measure, which provides information about the change in normal direction along the surface.

#### SD calculation

SD measures the depth of sulci and was calculated as the Euclidean distance between the central surface and its convex hull based on the spherical harmonic reconstructions.

### Construction of individual morphological similarity networks

A network is made up of nodes and edges between the nodes. In this study, we constructed large-scale morphological similarity networks with nodes denoting brain regions and edges denoting interregional similarity in intraregional distributions of morphological indices.

#### Definition of network nodes

To define network nodes, we employed two widely used surface atlases, the a2009s atlas (Destrieux et al., 2010) and the DK40 atlas (termed a2005s atlas in this study) (Desikan et al., 2006), which divided the cerebral cortex into 148 and 68 regions of interest (ROIs), respectively.

#### Definition of network edges

To estimate interregional morphological similarity, we utilized two measures, the *KLDs* and its variant, the *JSDs*, to estimate the similarity in intraregional distribution of morphological indices between regions. In mathematical statistics, the *KLD* is a measure of how one probability distribution is different from a second, reference probability distribution (Kullback and Leibler, 1951). This measure has been widely used in the fields of image processing and machine learning. First, we extracted values of all vertices within each ROI for each morphological index. Then, a probability density estimate was computed for each ROI and each morphological index based on a normal kernel function (MATLAB function, ksdensity). The resultant probability density functions (four per region) were further converted to probability distribution functions (PDFs). The *KLDs* between two PDFs *P* and *Q* is computed as:

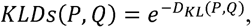

where *e* is the natural base and 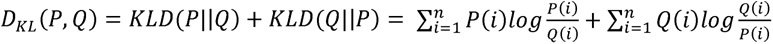, with *n* being the number of sample points (2^8^ in this study) (Wang et al., 2016). For the *JSDs* between *P* and *Q*, the formula is:

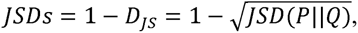

where 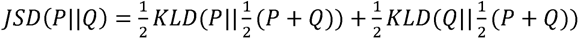. The value range for both *KLDs* and *JSDs* is [0, 1], with 0 and 1 denoting that two PDFs are absolutely different or exactly the same, respectively.

### Network analysis

Prior to graph-based topological characterization of morphological similarity networks derived above, a thresholding procedure was used to convert each network to a series of binary graphs. All topological analyses were performed with the GRETNA toolbox (Wang et al., 2015).

#### Threshold selection

Consistent with our previous study (Wang et al., 2016), we employed a sparsity-based thresholding procedure, where sparsity was defined as the ratio of the number of actual edges divided by the maximum possible number of edges in a graph. Given the lack of a conclusive method for selecting a single sparsity, a consecutive sparsity range was used in this study: [0.034 0.4] for the a2009s atlas and [0.063 0.4] for the a2005s atlas (interval = 0.02). The lower limits of the sparsity ranges were chosen to ensure that the resultant graphs would be estimable for the small-world attributes (Watts and Strogatz, 1998), that is, the average nodal degree (nodal degree is defined as the number of edges linked to a node) over all nodes of each thresholded graph is larger than 2 × log(*N*), with *N* denoting the number of nodes (i.e., 68 or 148 in this study). This criterion guarantees that the generated random networks are connected and thus the small world parameters can be successfully calculated (Watts and Strogatz, 1998). For the upper limits of the sparsity ranges, they were determined to guarantee that the thresholded graphs have sparse properties (He et al., 2007; Wang et al., 2009). All subsequent topological analyses were conducted at each sparsity level, and thus each network measure calculated below is a function or curve of sparsity. To provide sparsity-independent summary scalars, we computed the area under the curve (AUC; i.e., the integral over the entire sparsity range) for each network measure (Zhang et al., 2011), which were used to simplify the TRT reliability and statistical analyses.

#### Global and nodal network measures

For each graph, we calculated both global (clustering coefficient, *C*_p_; characteristic path length, *L*_p_; local efficiency, *E*_loc_; global efficiency, *E*_glob_; and modularity, *Q*) and nodal (nodal degree, *k*_i_; nodal efficiency, *e*_i_; and nodal betweenness, *b*_i_) measures. Detailed formulas and interpretations of these measures can be found elsewhere (Rubinov and Sporns, 2010; Wang et al., 2011) and are summarized in Table S1. To test whether the morphological similarity networks were non-randomly organized, all global measures were further normalized by dividing them by corresponding measures averaged over 100 matched random networks. The random networks were generated using a topological rewiring method (Maslov and Sneppen, 2002), which guaranteed the same degree distributions between real and random networks. Typically, a small-world, highly efficient and modular network should fulfill the following conditions: normalized *C*_p_ > 1 and normalized *L*_p_ ∼ 1, normalized *E*_loc_ > 1 and normalized *E*_glob_ ∼ 1 and normalized *Q* > 1.

### TRT reliability

We calculated the intraclass correlation coefficient (*ICC*) (Shrout and Fleiss, 1979) to quantify TRT reliability of morphological similarity networks. Formally, for a given measure repeatedly observed *k* times, the *ICC* was calculated as:

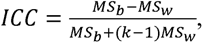

where *MS_b_* is the between-subject sum of squares and *MS_w_* is the within-subject sum of squares. ICC is close to 1 for reliable measures and 0 (negative) otherwise. In accordance with our previous studies (Wang et al., 2016; Wang et al., 2011), the TRT reliability scores were categorized as poor (*ICC* < 0.25), low (0.25 < *ICC <* 0.4), fair (0.4 < *ICC <* 0.6), good (0.6 < *ICC* < 0.75) and excellent (0.75 < *ICC* < 1).

In this study, the *ICC* was calculated for each interregional morphological similarity and each network measure (global and nodal) under all possible combinations of 4 morphological indices, 2 atlases and 2 similarity measures. This resulted in a total of 16 *ICC* matrices (148 × 148 or 68 × 68) for interregional morphological similarity, 160 *ICC* values for global network measures and 48 *ICC* vectors (148 or 68) for nodal network measures.

### Statistical analysis

#### Effects of different analytical strategies on morphological similarity networks

We explored effects of different choices of morphological index, brain parcellation atlas and similarity measure on morphological similarity networks at multiple levels. First, at the level of interregional morphological similarity, we first examined differences in the mean morphological similarity over all possible pairs of regions between the two parcellation atlases regardless of morphological indices and similarity measures (t-test). Then, for each brain parcellation atlas, we performed two-way repeated ANOVA on the morphological similarity between any pair of regions (10,878 ANOVA under the a2009s parcellation atlas and 2,278 ANOVA under the a2005s parcellation atlas). Second, at the level of global network organization, we performed three-way repeated ANOVA on each global network measure (10 ANOVA). Finally, at the level of local network organization, we first compared differences in the mean nodal centrality across all regions for each nodal centrality measure between the two parcellation atlases regardless of morphological indices and similarity measures (2 t-tests; nodal degree was excluded from this analysis because the mean nodal degree was equal among all participants in terms of the sparsity-based thresholding procedure). Then, under each brain parcellation atlas, we performed two-way repeated ANOVA on each nodal measure of each region (444 ANOVA under the a2009s parcellation atlas and 204 ANOVA under the a2005s parcellation atlas). For significant effects, simple effects were further examined (paired t-test). All results were corrected for multiple comparisons using the false discovery rate (FDR) procedure at the level of q < 0.05, when applicable.

#### Effects of different analytical strategies on TRT reliability of morphological similarity networks

Effects of different analytical strategies on the TRT reliability of morphological similarity networks were also examined at multiple levels. First, at the level of interregional morphological similarity, we first tested differences in the mean connectional *ICC* over all possible pairs of regions between the two parcellation atlases regardless of morphological indices and similarity measures (t-test). Then, under each brain parcellation atlas, we performed two-way repeated ANOVA on all connectional *ICC* values (2 ANOVA). Second, at the level of global network organization, we performed three-way repeated ANOVA on *ICC* values for all global network measures. Finally, at the level of local network organization, we first examined differences in the mean nodal *ICC* over all regions between the two parcellation atlases regardless of morphological indices and similarity measures (3 t-tests). Then, under each brain parcellation atlas, we performed two-way repeated ANOVA on all nodal *ICC* values for each nodal measure (6 ANOVA). For significant effects, simple effects were further examined (paired t-test). All results were corrected for multiple comparisons using the FDR procedure at the level of q < 0.05, when applicable.

In addition, we utilized Z-tests (McGraw and Wong, 1996) to locate edges, global measures and regions, whose *ICC* values were significantly affected by different analytical strategies. Specifically, for two given *ICC* values *ICC*_1_ and *ICC*_2_, we first transformed the difference between *ICC*_1_ and *ICC*_2_ into z:

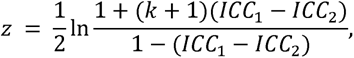

where *k* is the number of repeated observations (2 in this study). The *z* statistic has a mean of 0 with variance (McGraw and Wong, 1996):

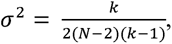

where *N* is the number of participants. Then, we transformed the *z* into a *Z* score with standard normal distribution:

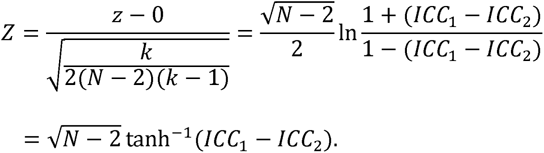

The Z-test was performed for each interregional morphological similarity and each nodal centrality measure for each region (FD/GI/SD/CT: *KLDs* versus *JSDs*; *KLDs*/*JSDs*: FD versus GI, FD versus SD, FD versus CT, GI versus SD, GI versus CT and SD versus CT), and each global network measure (FD/GI/SD/CT - a2005s/a2009s: *KLDs* versus *JSDs*; FD/GI/SD/CT - *KLDs*/*JSDs*: a2005s versus a2009s; a2005s/a2009s - *KLDs*/*JSDs*: FD versus GI, FD versus SD, FD versus CT, GI versus SD, GI versus CT and SD versus CT). All results were corrected for multiple comparisons using the FDR procedure at the level of q < 0.05, when applicable.

### Effects of sample size on morphological similarity networks

To test effects of different sample sizes on morphological similarity networks, we correlated the cross-subject mean nodal centrality map (concatenated across the three nodal measures) from a subgroup of participants with that from all participants. The subgroup of participants was randomly selected from all participants with sample size varying from 10 to 870 (interval = 10). This subgroup sampling procedure was repeated 1,000 times to generate 1,000 correlation coefficients at any fixed sample size, whose means and standard errors were calculated.

### Validation analysis

#### Twin subjects

The HCP S900 dataset includes twin subjects, which may yield bias for our results. Thus, we re-analyzed the HCP S900 data to estimate the reproducibility of our results by excluding twin subjects (338).

#### High-resolution parcellation atlas

In this study, we utilized the a2005s atlas and a2009s atlas for network node definition because they are two of the most commonly used surface parcelllation schemes in previous brain network studies (e.g., Buchanan et al., 2020; Rodríguez-Cruces et al., 2020; Seibert et al., 2011; Zhang et al., 2019). However, these two atlases are relatively coarser (68 and 148 regions, respectively), which may be insufficient to represent local characteristics of finer-level brain regions. Thus, we further re-analyzed the data by constructing individual morphological similarity networks with a high-resolution atlas, which divides the cerebral surface into 360 regions (termed MMP atlas; Glasser et al., 2016). It should be noted that we mainly focused on a more practical question in guiding studies of morphological similarity networks: whether brain parcellation atlases with higher resolutions will give rise to higher test-retest reliabilities. We did not re-analyzed the HCP S900 dataset because of the huge amount of computation, exponentially increased computing time with the number of network nodes and a growing body of evidence for parcellation-dependent human brain networks (Arslan et al., 2018).

##### Smoothing size of CT maps

In this study, individual CT maps were smoothed using a Gaussian kernel with 15-mm full width at half maximum, which was smaller than those for the other three morphological indices (Gaussian kernel with 25-mm full width at half maximum). To test whether the differences in smoothing size could lead to relatively poor performance in TRT reliability for CT-based morphological similarity networks (see Results), we smoothed individual CT maps again (BNU TRT dataset) using a Gaussian kernel with 25-mm full width at half maximum, followed by network parameter and ICC calculation.

##### Brain size and regional size

Brain size is an important confounding factor for analysis of brain morphology. In addition, previous functional (Wang et al., 2009) and morphological (Seidlitz et al., 2018) brain network studies showed the existence of relationships between regional size and nodal centrality. Thus, in this study we calculated cross-subject Pearson correlation between each global network measure and global morphological values, and cross-node, person-level Pearson correlation between each nodal centrality measure and regional size (defined as the number of vertex in a region) for each type of morphological similarity networks.

## Results

### Interregional similarities of morphological similarity networks

#### Similarity matrices

Unless stated otherwise, all results reported are exemplified using the a2009s atlas for brain parcellation and the *JSD*s for interregional similarity estimation.

Figure 2 shows the mean morphological similarity matrices derived from the four morphological indices of FD, GI, SD and CT. In general, the cross-subject mean interregional similarity was large with relatively small variance for each edge of each type of morphological similarity networks (FD: 0.725 ± 0.074; GI: 0.724 ± 0.069; SD: 0.717 ± 0.121; CT: 0.737 ± 0.072). Nonetheless, morphological index-dependent similarity patterns were evident. For example, the SD-based morphological similarity networks were clearly different from the others by visual inspection. This was further confirmed by the relatively low Spearman’s rank correlations in the mean similarity matrix between any pair of morphological similarity networks (*r*_FD-GI_ = 0.213; *r*_FD-SD_ = 0.136; *r*_FD-CT_ = 0.156; *r*_GI-SD_ = 0.156; *r*_GI-CT_ = 0.304; *r*_SD-CT_ = 0.289). Interestingly, we consistently found that the mean interregional similarity for edges linking geometrically corresponding regions between two hemispheres (i.e., homotopic connections) was significantly higher than that for edges linking nonhomotopic regions (i.e., heterotopic connections) regardless of the types of morphological similarity networks (t-test; FD: *P ≈* 0; GI: *P ≈* 0; SD: *P ≈* 0; CT: *P ≈* 0).

**Figure 2.**
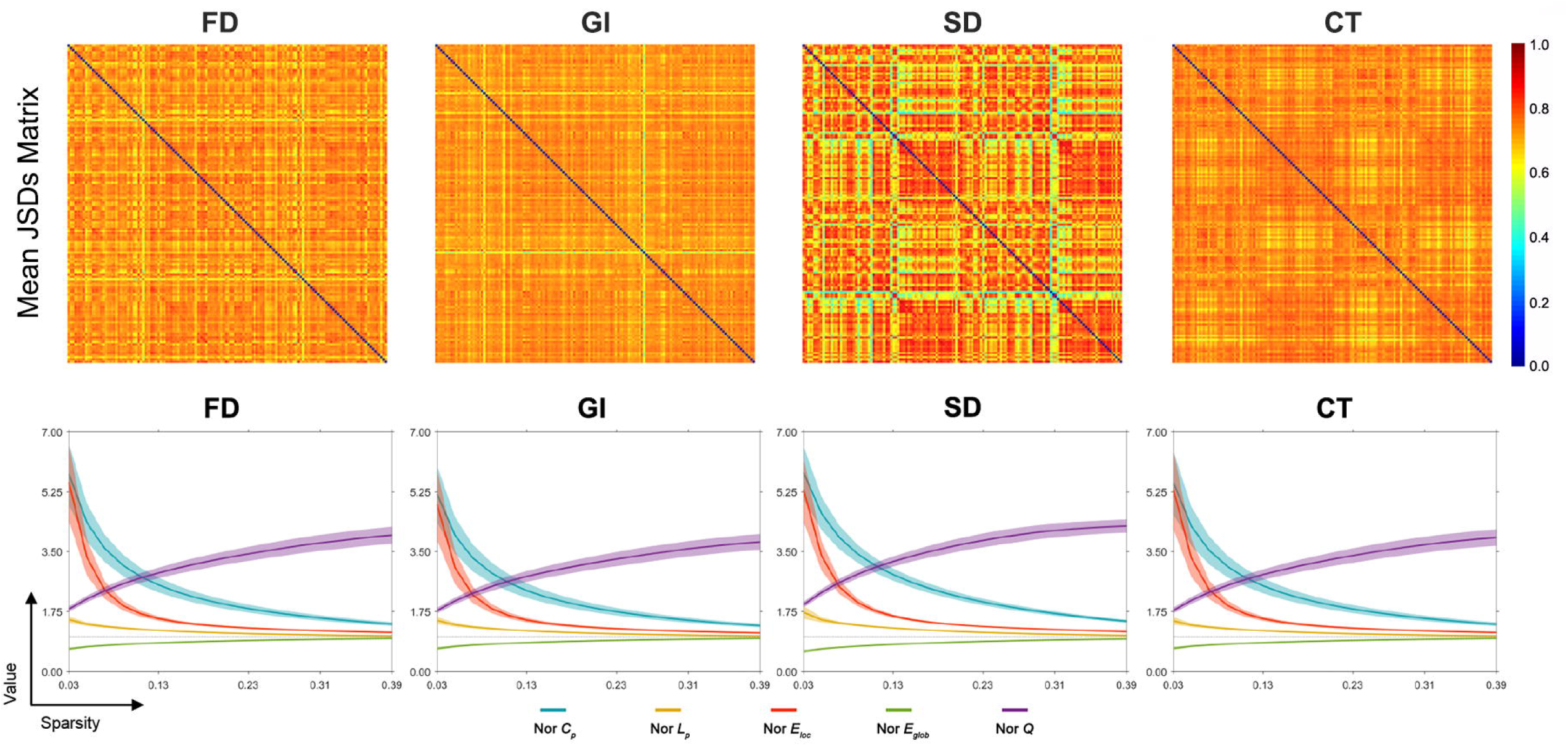
Mean similarity matrices and global network measures of morphological similarity networks. Upper panel: mean similarity matrices of morphological similarity networks (brain parcellation: a2009s; similarity estimation: *JSDs*). Although the similarity strength was large for each connection regardless of the choice of morphological index, differential similarity patterns were evident. Lower panel: global organization of morphological similarity networks (brain parcellation: a2009s; similarity estimation: *JSDs*). Compared with matched random networks, individual morphological similarity networks exhibited higher clustering coefficient (i.e., normalized *C*_p_ > 1) and approximately equal characteristic path length (i.e., normalized *L*_p_ ∼ 1), higher local efficiency (i.e., normalized *E*_loc_ > 1) and approximately equal global efficiency (i.e., normalized *E*_glob_ ∼ 1), and higher modularity (i.e., normalized *Q* > 1), indicating small-world, highly efficient and modular organization. FD, fractal dimension; GI, gyrification index; SD, sulcal depth; CT, cortical thickness.

Both descriptive and inferential statistical results mentioned above were consistently observed regardless choices of brain parcellation atlas and similarity measure.

#### Effects of morphological index, brain parcellation and similarity measure

Significant differences were observed in the mean interregional morphological similarity across all edges when using either the a2009s or the a2005s atlases for brain parcellation (*P ≈* 0). Further analyses of the interregional morphological similarity of each edge under both atlases consistently revealed that all edges were significantly affected by choices of at least one factor of morphological index or similarity measure (*P* < 0.05, FDR corrected). These findings jointly suggest widespread influence of different analytical strategies on interregional similarity patterns of morphological similarity networks. Notably, consistent with our previous study (Wang et al., 2016), no post hoc analyses were conducted since comparisons of numerical values have no utility with respect to the selection of optimal analytical strategies.

### Global organization of morphological similarity networks

#### Small-worldness, efficiency and modularity

Each individual morphological similarity network exhibited small-world organization, high parallel efficiency and modular structure over the entire sparsity range regardless of the morphological indices. That is, compared with matched random networks, individual morphological similarity networks had higher *C*_p_ and approximately equal *L*_p_, higher *E*_loc_ and approximately equal *E*_glob_, and higher *Q* (Fig. 2). These organizational principles were consistently observed under other combinations of brain parcellation atlases and similarity measures.

#### Effects of morphological index, brain parcellation and similarity measure

Despite the common organizational principles, three-way repeated ANOVA revealed that quantitative values of all global network measures were significantly affected by choices of morphological index, brain parcellation atlas and/or similarity measure (*P* < 0.05, FDR corrected). Again, no post hoc analyses were further conducted, as explained above.

### Local organization of morphological similarity networks

#### Hubs

Visual inspection indicated that the overall patterns of nodal centrality measures were spatially heterogeneous and depended on different analytical strategies. Thus, we first calculated Pearson correlation coefficients between each pair of nodal centrality maps (4 morphological indices × 2 similarity measures × 3 nodal centrality measures) averaged across all participants under each parcellation atlas. The significance level for each correlation coefficient was estimated using a recently proposed approach that corrects for spatial autocorrelation of brain maps (Burt et al., 2020). Specifically, for each pair of nodal centrality maps used in the correlation analysis, one of them was randomly selected as a target map and 10,000 surrogate maps were generated that matched with the target map with respect to spatial autocorrelation. Each of the surrogate maps was then used to re-compute the Pearson correlation coefficient with the other map, yielding a null distribution for the expected value of Pearson correlation by chance. Subsequently, a *P*-value was estimated as the fraction of surrogate maps which generated a Pearson correlation equal to or greater than the real Pearson correlation. Of note, variability into the estimate of the *P*-value introduced by finite sampling size was accounted for by using a binomial distribution to estimate the size of sampling fluctuations. As shown in Figure 3, we found that under each parcellation atlas different nodal centrality measures exhibited highly similar spatial patterns regardless of the choices of similarity measure (a2009s: *r* = 0.891 ± 0.087; a2005s: *r* = 0.714 ± 0.214), while the spatial similarities became dramatically low when the correlations were calculated between different morphological indices (a2009s: *r* = 0.306 ± 0.053; a2005s: *r* = 0.312 ± 0.134). These findings indicate that the factor of morphological index dominates spatial distributions of nodal centrality measures. Notably, the correlation pattern in regional centrality profiles was obviously different from the correlation pattern in regional mean morphological values (Table S2). For example, compared with the moderate negative correlations in regional mean values between GI and CT (a2009s: *r* = −0.476 ± 0.083; a2005s: *r* = −0.501 ± 0.083), low positive correlations were observed in regional centrality profiles between GI-based and CT-based morphological similarity networks (a2009s/*JSDs*: *r* = 0.363 ± 0.048; a2009s/*KLDs*: *r* = 0.366 ± 0.047; a2005s/JSDs: *r* = 0.276 ± 0.087; a2005s/KLDs: *r* = 0.278 ± 0.087). The discrepancies suggest that our findings are not affected by the existence of some spatial correlations in regional mean values between different morphological indices.

**Figure 3.**
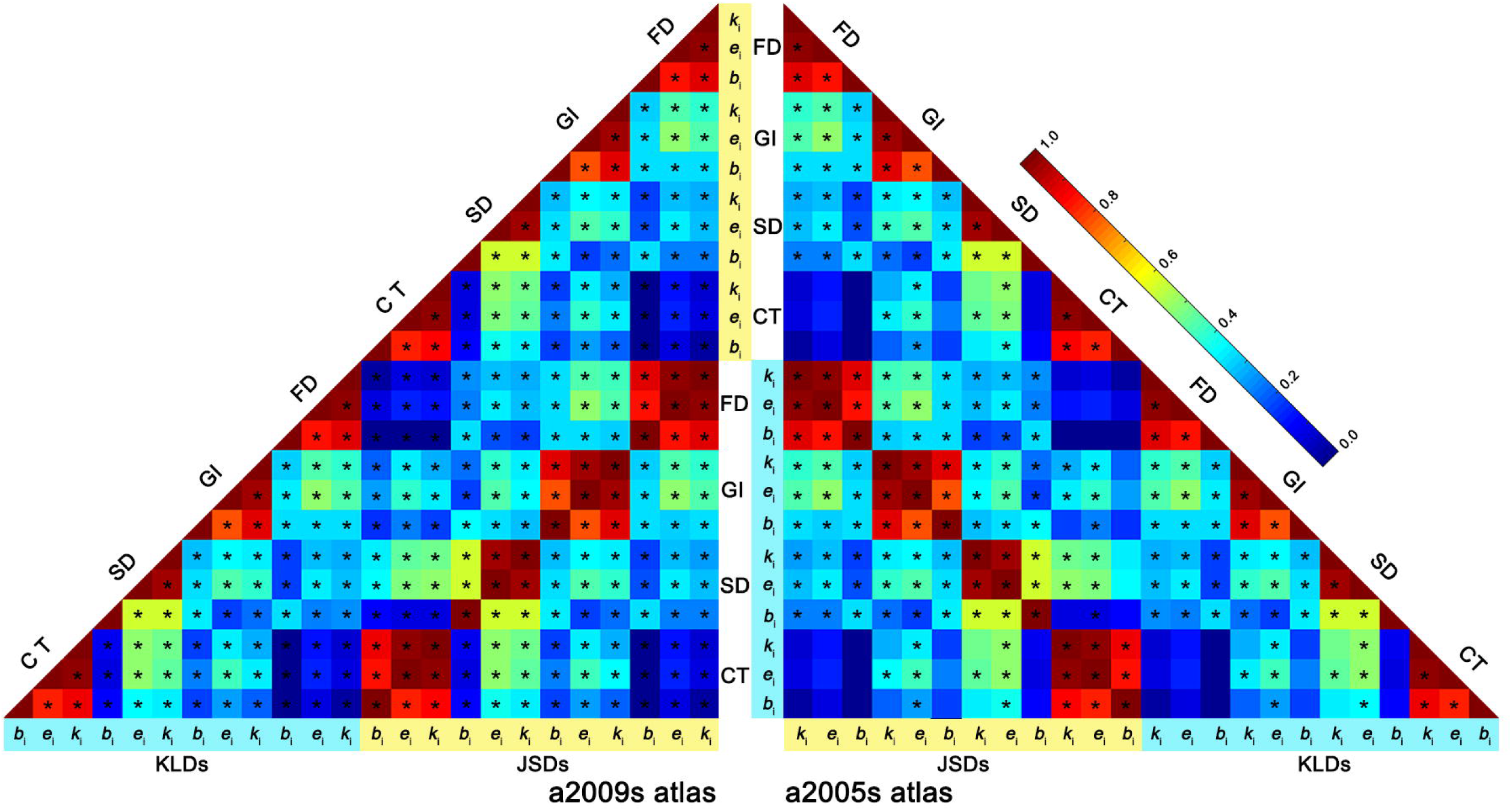
Spatial correlations of mean nodal centrality maps under different analytical strategies. Under each parcellation atlas, high correlations were observed between different nodal metrics and between different similarity measures; however, dramatically low correlations were observed between different morphological indices. *JSDs*, Jensen-Shannon divergence-based similarity; *KLDs*, Kullback-Leibler divergence-based similarity; FD, fractal dimension; GI, gyrification index; SD, sulcal depth; CT, cortical thickness; *k*_i_, nodal degree; *e*_i_, nodal efficiency; *b*_i_, nodal betweenness; *, *P* < 0.05, Bonferroni corrected.

Furthermore, given central roles of highly connected regions (i.e., hubs), we identified hubs, which were defined as regions with values in the top 10% of each nodal centrality measure. We found several common features of the identified hubs, despite differential spatial distributions across different morphological indices and nodal centrality measures (Fig.S1). First, the majority of hubs was located at bilaterally homologous regions, with the ratio of such hubs to all hubs varying between 40.0% and 80.0% (mean = 62.2%) among different morphological indices and nodal centrality measures. Second, most hubs were located at brain sulci, especially at the junction between different lobes, with the ratio of such hubs to all hubs varying between 60.0% and 86.7% (mean = 75.0%) among different morphological indices and nodal measures. Finally, several regions were consistently identified hubs that were independent of choices of morphological index and nodal centrality measure. These features also existed under other combinations of brain parcellation atlases and similarity measures (except for the second feature when regional parcellation was based on the a2005s atlas, which was composed of only gyral-based regions).

Finally, we defined a consistent hub score (*CHs*), which was calculated as the number of times a region was identified as a hub under all combinations of morphological index, similarity measure and nodal centrality measure under each parcellation atlas. Figure 4 shows regions with the top 10% highest values of *CHs*.

**Figure 4.**
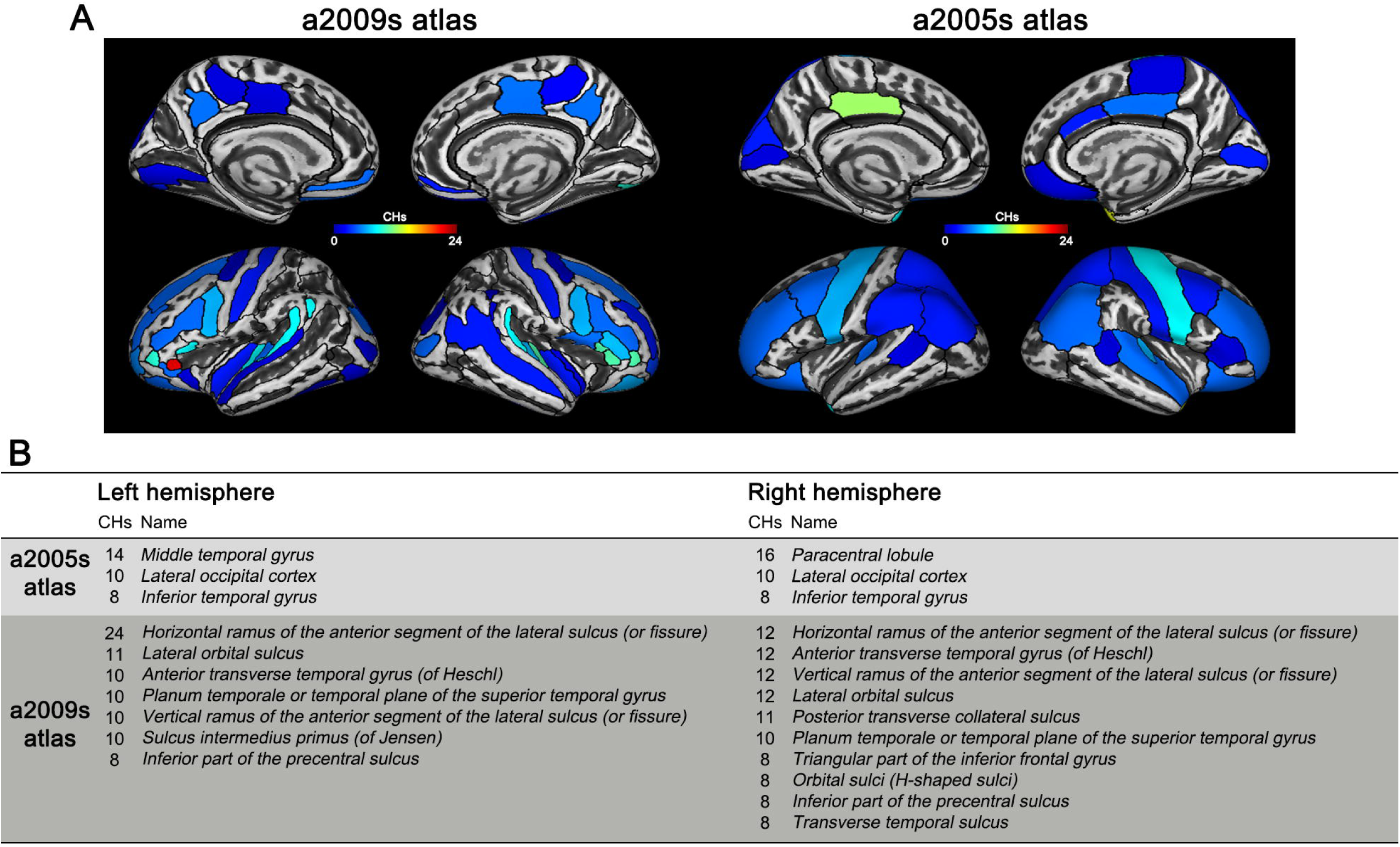
Consistent hubs under different analytical strategies. A specific set of regions were consistently identified as hubs, among which regions with the top 10% of *CHs* were listed under each parcellation atlas. *CHs*, consistent hub scores.

#### Effects of morphological index, brain parcellation and similarity measure

Between-atlas comparisons of the mean centrality value across all regions revealed significant differences in nodal efficiency and betweenness (*P ≈* 0). Further two-way repeated ANOVA revealed that all nodal centrality measures of all regions were significantly affected by choices of morphological index and/or similarity measure under both the a2009s and a2005s atlases (*P* < 0.05, FDR corrected).

### TRT reliability of interregional similarities of morphological similarity networks

#### ICC of similarity matrices

Based on the BNU TRT dataset, extremely high correlations across edges were observed in the mean similarity matrices between the two sessions regardless of different morphological indices (all *r* > 0.966, Fig. S2). Further TRT reliability analysis on the interregional similarity of each edge revealed that 99.8%, 88.8%, 96.6% and 80.6% of all edges exhibited good or above TRT reliability (i.e., *ICC* > 0.6) for the FD-, GI-, SD- and CT-based morphological similarity networks, respectively. The high TRT reliability was also observed under other combinations of brain parcellation atlases and similarity measures.

#### Effects of morphological index, brain parcellation and similarity measure

First, between-atlas comparison of all connectional *ICC* values (concatenated across different morphological indices and similarity measures) revealed that the TRT reliabilities were significantly higher for the a2009s than a2005s atlas (*ICC*_a2009s_ = 0.801 ± 0.135, *ICC*_a2005s_ = 0.782 ± 0.149; *t*_102902_ = 17.378, *P* < 0.001). Then, under each brain parcellation atlas, two-way repeated ANOVA consistently revealed that the mean *ICC* values across all edges were significantly modulated by choices of morphological index and similarity measure with a significant interaction (all *P ≈* 0). Post hoc analyses revealed that: 1) the *ICC* differed significantly among brain networks constructed using different morphological indices no matter the choices of brain parcellation atlas and similarity measure (FD > SD > GI > CT; all *P* < 0.05, FDR corrected across 24 t-tests in total); and 2) the *ICC* differed significantly among morphological similarity networks constructed using different similarity measures regardless of choices of brain parcellation atlases (*JSDs* > *KLDs* for FD-, GI- and SD-based morphological similarity networks, and *KLDs* > *JSDs* for CT-based morphological similarity networks; all *P* < 0.05, FDR corrected across 8 t-tests in total). Finally, we located edges whose TRT reliabilities were affected by choices of morphological index and similarity measure. The results showed that: 1) no edges exhibited significant differences in TRT reliabilities when morphological similarity networks were constructed using *JSDs* or *KLDs* for interregional similarity estimation (*P* > 0.05, FDR corrected); and 2) the TRT reliabilities of up to 7.3% of the edges for the a2009s atlas and of up to 19.2% of the edges for the a2005s atlas were significantly different when morphological similarity networks were constructed using different morphological indices (*P* < 0.05, FDR corrected). Interestingly, the TRT reliabilities of the identified edges exhibited a common pattern of FD > SD > GI > CT (Fig. S3).

### TRT reliability of global network measures of morphological similarity networks

#### ICC of global network measures

Individual morphological similarity networks exhibited small-world organization, high parallel efficiency and modular structure regardless of the data sessions (Fig. S4). The *ICC* analysis further revealed that most global network measures exhibited fair to excellent TRT reliabilities (Fig. S5).

#### Effects of morphological index, brain parcellation and similarity measure

Three-way repeated ANOVA revealed that the *ICC* of global measures were significantly modulated by the factors of morphological index (*F*_3,159_ = 105.728, *P ≈* 0) and similarity measure (*F*_1,159_ = 5.296, *P* = 0.047), and the modulation depended on choices of brain parcellation atlas (*F*_3,159_ = 8.349, *P* < 0.001 for the interaction between morphological index and brain parcellation atlas, and *F*_1,159_ = 15.351, *P* = 0.004 for the interaction between similarity measure and brain parcellation atlas) (Fig. 5). Further post hoc analyses revealed that: 1) the TRT reliabilities of global network measures were significantly higher for *JSDs*-based than for *KLDs*-based morphological similarity networks under the a2005s (*t*_39_ = 2.834, *P* = 0.007) but not the a2009s (*t*_39_ = 0.093, *P* = 0.926) atlas; and 2) the TRT reliabilities of global network measures differed significantly between any pair of morphological indices with a pattern of SD > FD > CT > GI under the a2005s atlas (all *P* < 0.05, FDR corrected), while they were significantly higher for the FD- and SD- than for the GI- and CT-based morphological similarity networks under the a2009s atlas (*P* < 0.05, FDR corrected). Finally, we identified specific global measures whose TRT reliabilities were affected by choices of morphological index, brain parcellation atlas and similarity measure. The results showed that: 1) no global measures exhibited significant differences in TRT reliabilities when morphological similarity networks were constructed using the a2009s or a2005s atlas for brain parcellation (*P* > 0.05, FDR corrected); 2) no global measures exhibited significant differences in TRT reliabilities when morphological similarity networks were constructed using *JSDs* or *KLDs* for interregional similarity estimation (*P* > 0.05, FDR corrected); and 3) the TRT reliabilities of many global measures were significantly affected by choices of morphological index (*P* > 0.05, FDR corrected) with a common pattern of FD- and SD- > GI- and CT-based morphological similarity networks (Fig. S6).

**Figure 5.**
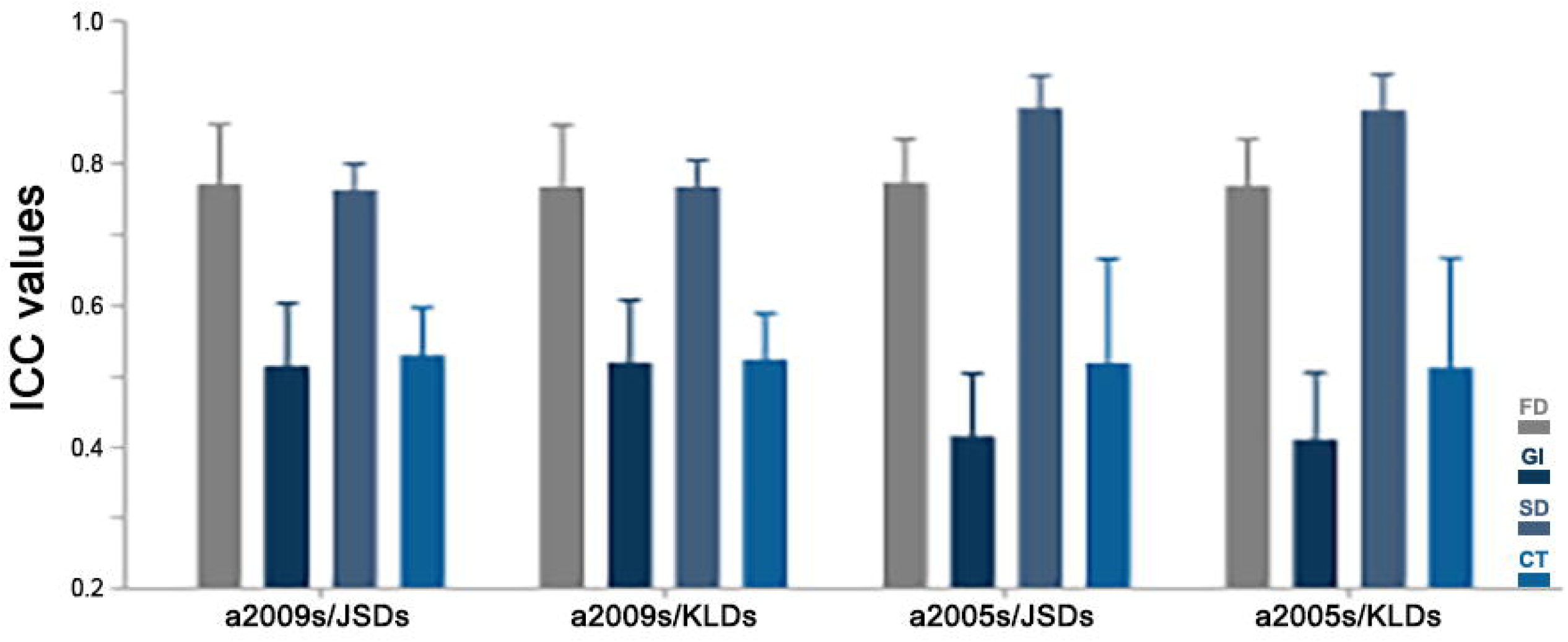
TRT reliability values for global metrics under different analytical strategies. TRT reliability of global network metrics was significantly affected by choices of morphological index, brain parcellation atlas and similarity measure, with a general pattern of FD and SD > GI and CT and *JSDs* > *KLDs*. *ICC*, intraclass correlation; *JSDs*, Jensen-Shannon divergence-based similarity; *KLDs*, Kullback-Leibler divergence-based similarity; FD, fractal dimension; GI, gyrification index; SD, sulcal depth; CT, cortical thickness.

### TRT reliability of nodal network measures of morphological similarity networks

#### ICC of nodal centralities

Similar to nodal centrality maps, the overall patterns of nodal *ICC* were spatially heterogeneous and depended on different analytical strategies. Thus, we also calculated the Pearson correlation between any pair of *ICC* maps under each atlas (4 morphological indices × 2 similarity measures × 3 nodal measures). Again, the significance levels of the resultant correlation coefficients were estimated using the approach from (Burt et al., 2020) to correct for spatial autocorrelation of brain maps. The results showed high correlation coefficients between different nodal centrality measures regardless of the choice of similarity measure (a2009s: *r* = 0.869 ± 0.139; a2005s: *r* = 0.574 ± 0.326). However, the correlation coefficients between different morphological indices were dramatically low (a2009s: *r* = 0274 ± 0.145; a2005s: *r* = 0.258 ± 0.120) (Fig. S7).

We further divided all brain regions into five categories in terms of their *ICC* values. As shown in Figures 6 and S8, a considerable proportion of brain regions exhibited good to excellent TRT reliabilities for most morphological indices and nodal centrality measures. Nevertheless, it was evident that the proportions were higher for the FD- and SD- than for the GI- and CT-based morphological similarity networks and were higher for nodal degree and efficiency than for nodal betweenness.

**Figure 6.**
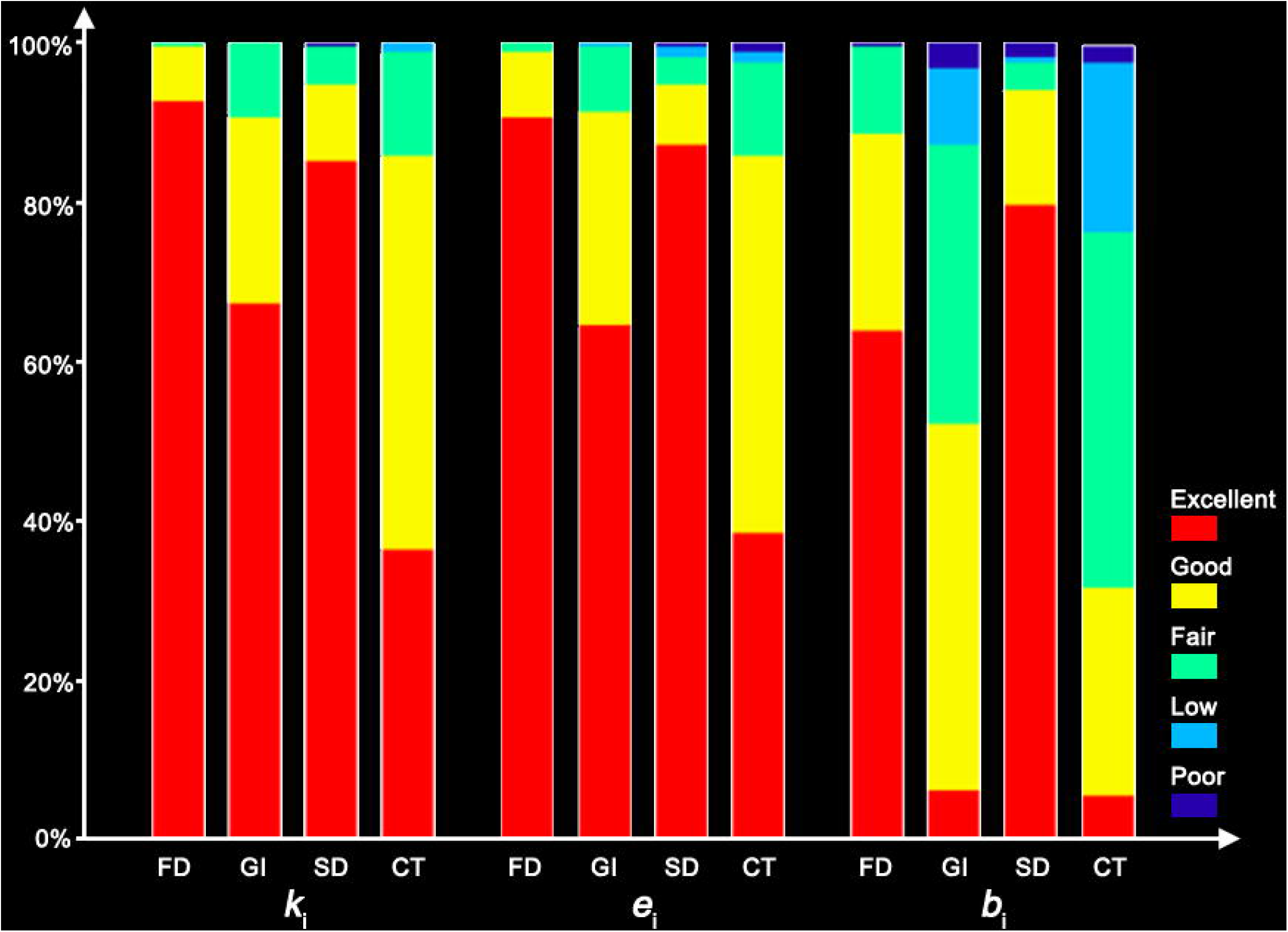
Proportions of regions at different levels of nodal TRT reliability (brain parcellation: a2009s; similarity estimation: *JSDs*). The proportions of regions showing excellent TRT reliability were significantly higher for nodal degree and efficiency than for nodal betweenness, and for FD- and SD-based than for GI- and CT-based morphological similarity networks. FD, fractal dimension; GI, gyrification index; SD, sulcal depth; CT, cortical thickness; *k*_i_, nodal degree; *e*_i_, nodal efficiency; *b*_i_, nodal betweenness.

Finally, we identified regions under each brain parcellation atlas that consistently exhibited high TRT reliabilities regardless of different analytical strategies by defining a consistent excellent reliability score (*CERs*) as the number of times a region exhibited excellent TRT reliability (*ICC* > 0.75) under all combinations of morphological indices, similarity measures and nodal metrics. Figure 7 shows the regions with the top 10% highest values of *CERs*.

**Figure 7.**
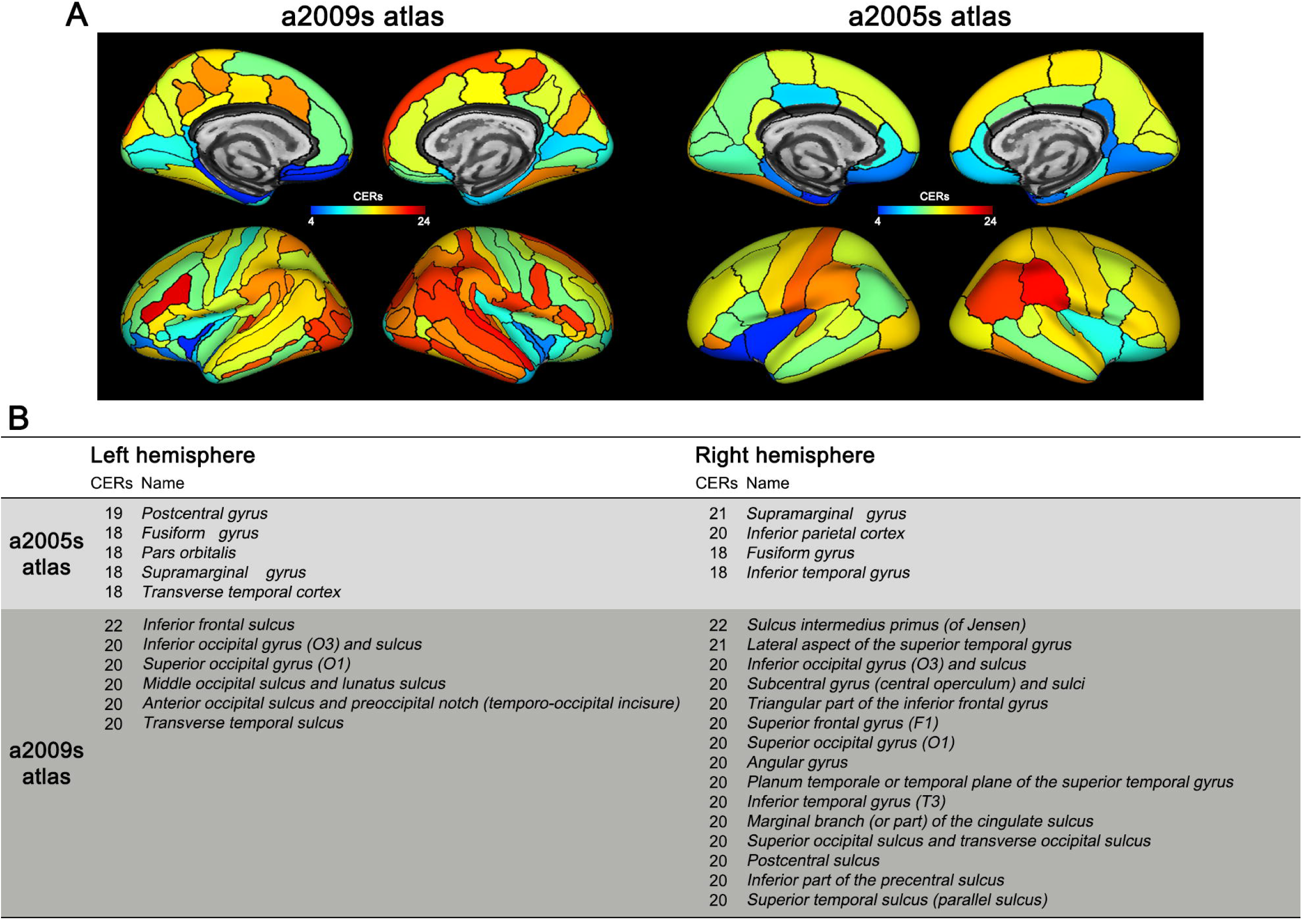
Regions consistently showing excellent TRT reliability under different analytical strategies. A specific set of regions were consistently identified to show excellent TRT reliability, among which regions with the top 10% of *CERs* are listed under each parcellation atlas. *CERs*, consistent excellent reliability scores.

#### Effects of morphological index, brain parcellation and similarity measure

First, comparison of all nodal *ICC* values (concatenated across different morphological indices and similarity measures) between the two parcellation atlases showed that the TRT reliabilities were significantly higher for the a2009s than for the a2005s atlas for each nodal measure (*k*_i_: *ICC*_a2009s_ = 0.803 ± 0.135, *ICC*_a2005s_ = 0.778 ± 0.149; *t_1710_* = 3.557, *P ≈* 0; *e*_i_: *ICC*_a2009s_ = 0.803 ± 0.136, *ICC*_a2005s_ = 0.773 ± 0.154; *t_1710_* = 4.081, *P ≈* 0; *b*_i_: *ICC*_a2009s_ = 0.665 ± 0.189, *ICC*_a2005s_ = 0.624 ± 0.207; *t_1710_* = 4.023, *P ≈* 0).

Then, under each brain parcellation atlas, two-way repeated ANOVA revealed that choices of morphological index and similarity measure significantly affected the mean *ICC* of nodal degree (all *P* < 0.004) and nodal efficiency (all *P* < 0.006) with nonsignificant interaction effects (all *P* > 0.05). For nodal betweenness, significant effects were only observed for the factor of morphological index under both the a2009s and a2005s atlases (both *P ≈* 0). Further post hoc analyses revealed that 1) the TRT reliabilities of nodal degree and nodal efficiency were higher for *JSDs*-based than *KLDs*-based morphological similarity networks under each brain parcellation atlas (all *P* < 0.05, FDR corrected across 4 t-tests in total); and 2) the TRT reliabilities of nodal degree and efficiency exhibited a pattern of FD- > SD- > GI- > CT-based morphological similarity networks under the a2009s atlas and a pattern of FD- and SD- > GI- > CT-based morphological similarity networks under the a2005s atlas; and nodal betweenness exhibited a pattern of SD- > FD- > GI- > CT-based morphological similarity networks for both atlases (all *P* < 0.05, FDR corrected across 36 t-tests in total).

Finally, we identified regions whose TRT reliabilities were affected by choices of morphological index and similarity measure. We found that: 1) no regions exhibited significant differences in TRT reliabilities for any nodal centrality measure when morphological similarity networks were constructed using *JSDs* or *KLDs* for interregional similarity estimation (*P* > 0.05, FDR corrected); and 2) the TRT reliabilities of up to 43.2% of the regions under the a2009s atlas and of up to 61.8% of the regions under the a2005s atlas were significantly different when morphological similarity networks were constructed using different morphological indices (*P* < 0.05, FDR corrected). Interestingly, the TRT reliabilities of the identified regions exhibited a common pattern of FD- and SD- > GI- and CT-based morphological similarity networks (Figs. S9-S12).

#### Differences in TRT reliability between hub and non-hub regions

We compared the TRT reliability between hub and non-hub regions under each analytical combination of morphological index, brain parcellation atlas and similarity measure (permutation test, 10,000 times). No significant differences were found regardless of the analytical strategies (*P* > 0.05, FDR corrected).

### Effects of sample size on individual morphological similarity networks

Highly positive correlations were observed between the nodal centrality profiles (concatenated across nodal measures) averaged over 10 randomly selected samples and the mean nodal centrality profile across all participants (*r*_FD_ = 0.944 ± 0.007; *r*_GI_ = 0.936 ± 0.007; *r*_SD_ = 0.970 ± 0.005; *r*_CT_ = 0.936 ± 0.008). With increasing sample size, the correlations continually increased and the variability naturally decreased, with a tipping point observed when samples exceeded ∼70 participants (*r*_FD_ = 0.992 ± 0.001; *r*_GI_ = 0.991 ± 0.001; *r*_SD_ = 0.996 ± 0.001; *r*_CT_ = 0.991 ± 0.001) (Fig. 8). More importantly, this feature was observed under other combinations of brain parcellation atlases and similarity measures.

**Figure 8.**
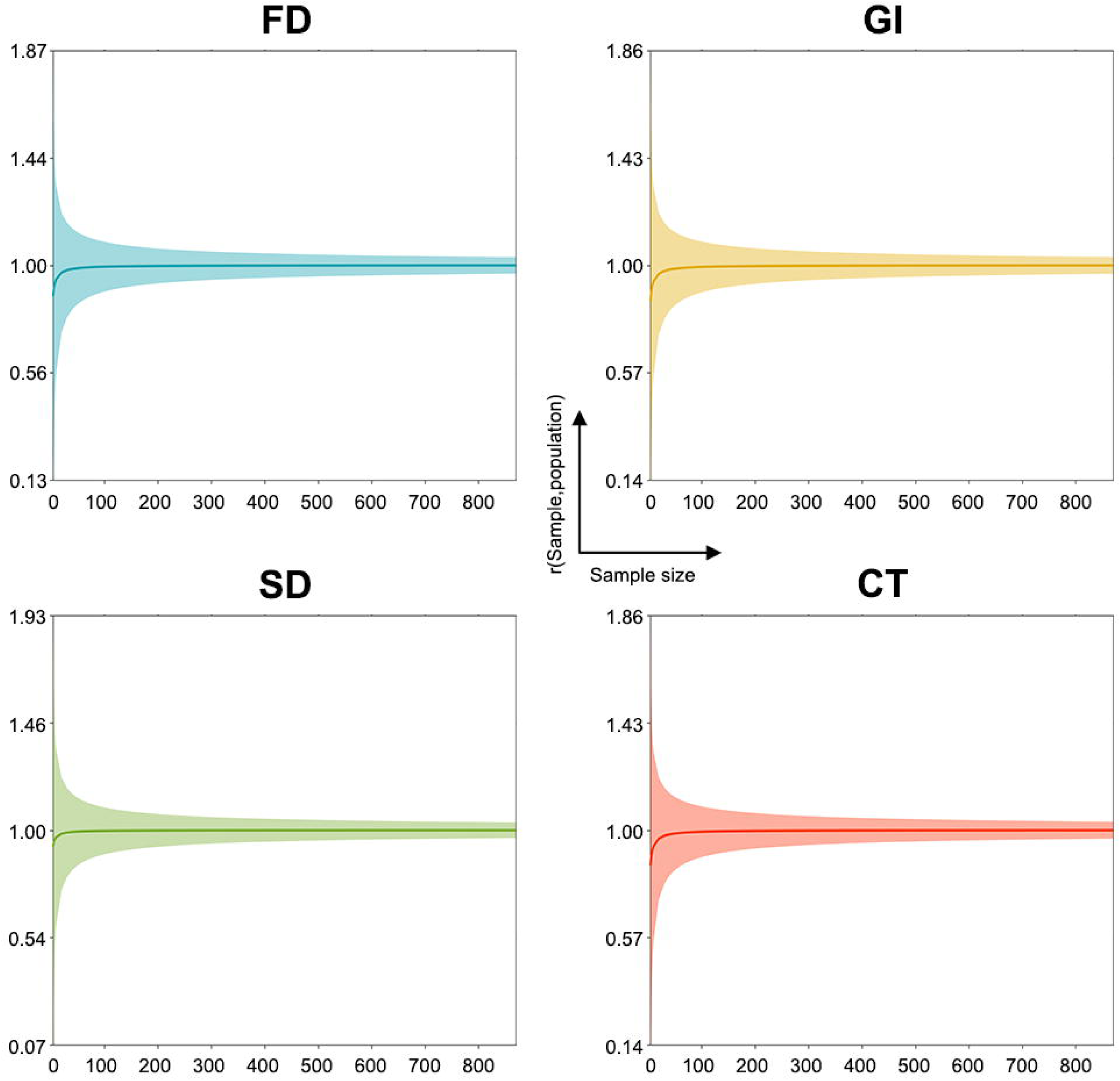
Effect of different sample sizes on morphological similarity networks (brain parcellation: a2009s; similarity estimation: *JSD*s). With increasing sample size, a continuous increase was observed in the Pearson correlation coefficients between nodal centrality profiles that were separately averaged for 1000 subgroups of participants and nodal centrality profiles that were averaged over all participants, while the variability naturally decreased, with a tipping point observed when samples exceeded ∼70 participants. FD, fractal dimension; GI, gyrification index; SD, sulcal depth; CT, cortical thickness.

### Reproducibility of our results

#### Effects of twin subjects

After excluding twin subjects from the HCP S900 dataset, we found that all results reported remained largely unchanged, indicating little effects of twin subjects on our results. The new results are briefly summarized below.

##### 1) Edge-level analysis

we found: a) relatively low Spearman’s rank correlations in the mean similarity matrix between any pair of morphological similarity networks (*r*_FD-GI_ = 0.240; *r*_FD-SD_ = 0.154; *r*_FD-CT_ = 0.170; *r*_GI-SD_ = 0.167; *r*_GI-CT_ = 0.304; *r*_SD-CT_ = 0.304); b) higher mean interregional similarity for edges linking homotopic regions than edges linking heterotopic regions regardless of the types of morphological similarity networks (FD: *t* = 13.833, *P ≈* 0; GI: *t* = 12.321, *P ≈* 0; SD: *t* = 20.844, *P ≈* 0; CT: *t* = 15.821, *P ≈* 0); c) significant differences in the mean interregional morphological similarity between the a2005s atlas and a2009s atlas (*t* = 91.119, *P ≈* 0); and d) significant effects of morphological index and/or similarity measure on the interregional morphological similarity of each edge under the two parcellation atlases (two-way repeated ANOVA; *P* < 0.05, FDR corrected).

##### 2) Global-level analysis

We found that all global network measures were significantly affected by choices of morphological index, brain parcellation atlas and/or similarity measure (three-way repeated ANOVA, *P* < 0.05, FDR corrected).

##### 3) Nodal-level analysis

We found: a) highly similar spatial patterns among different nodal centrality measures for the same morphological indices (a2009s: *r* = 0.869 ± 0.084; a2005s: *r* = 0.844 ± 0.139) but dramatically low spatial similarities between different morphological indices (a2009s: *r* = 0.305 ± 0.053; a2005s: *r* = 0.267 ± 0.144); b) significant differences in the mean centrality across all regions between the a2005s atlas and a2009s atlas (nodal efficiency: *t* = 230.836, *P ≈* 0; nodal betweenness: *t* = 560.849, *P ≈* 0); and c) significant effects of morphological index and/or similarity measure on each nodal centrality measure of each region under the two atlases (two-way repeated ANOVA; *P* < 0.05, FDR corrected).

##### 4) Analysis of sample size

We found that the correlation of mean nodal centrality profiles between randomly selected samples and all participants continually increased with increasing sample size, with a tipping point observed when samples exceeded ∼70 participants (*r*_FD_ = 0.992 ± 0.001; *r*_GI_ = 0.991 ± 0.001; *r*_SD_ = 0.996 ± 0.001; *r*_CT_ = 0.991 ± 0.001).

#### Effects of high-resolution parcellation atlas

Based on randomly selected 50 participants from the HCP S900 dataset, we found the existence of small-worldness, high parallel efficiency, modularity and hubs for surface-based single-subject morphological similarity networks. Moreover, these network measures derived from the MMP atlas were significantly different from those derived from the a2005s atlas and a2009s atlas, and were dependent on choices of morphological index and/or similarity measure. All these findings are as expected and in line with the main results.

Based on the BNU TRT dataset, we found that compared with the a2005s atlas and a2009s atlas, the MMP atlas was associated with significantly higher TRT reliabilities in interregional morphological similarities (a2005s: 0.782±0.149; a2009s: 0.801±0.135; MMP: 0.817±0.125), global (a2005s: 0.644±0.208; a2009s: 0.644±0.140; MMP: 0.673±0.130) and local (a2005s: 0.725±0.186; a2009s: 0.757±0.169; MMP: 0.773±0.152) network measures. It should be noted that this general conclusion does not always hold for global network measures when considering the other two factors of morphological index and similarity measure. All TRT reliability results in this study are summarized in Table 1.

**Table 1.** Summary of TRT reliability of morphological brain networks under different analytical strategies

| Parcellation atlas | Similarity measure | Morphological index | Interregional similarity | Global organization | Local organization |
| --- | --- | --- | --- | --- | --- |
| a2005s | *KLDs* | FD | 0.876±0.128 | 0.768±0.064 | 0.830±0.111 |
|  |  | GI | 0.734±0.141 | 0.413±0.093 | 0.663±0.166 |
|  |  | SD | 0.830±0.167 | 0.873±0.050 | 0.834±0.135 |
|  |  | CT | 0.636±0.161 | 0.514±0.152 | 0.572±0.174 |
|  | *JSDs* | FD | **0.877±0.127** | 0.772±0.060 | 0.829±0.112 |
|  |  | GI | 0.735±0.141 | 0.418±0.087 | 0.664±0.166 |
|  |  | SD | 0.842±0.153 | **0.877±0.045** | **0.835±0.135** |
|  |  | CT | 0.634±0.161 | 0.519±0.146 | 0.572±0.174 |
| a2009s | *KLDs* | FD | 0.883±0.098 | 0.766±0.086 | 0.842±0.115 |
|  |  | GI | 0.749±0.132 | 0.520±0.087 | 0.704±0.159 |
|  |  | SD | 0.853±0.150 | 0.766±0.037 | 0.840±0.145 |
|  |  | CT | 0.695±0.134 | 0.525±0.064 | 0.641±0.154 |
|  | *JSDs* | FD | **0.883±0.098** | **0.770±0.084** | **0.842±0.114** |
|  |  | GI | 0.749±0.133 | 0.516±0.087 | 0.705±0.158 |
|  |  | SD | 0.863±0.132 | 0.761±0.037 | 0.841±0.144 |
|  |  | CT | 0.693±0.136 | 0.531±0.067 | 0.641±0.155 |
| MMP | *KLDs* | FD | 0.892±0.077 | 0.743±0.073 | 0.852±0.101 |
|  |  | GI | 0.767±0.111 | 0.606±0.073 | 0.728±0.140 |
|  |  | SD | 0.886±0.121 | 0.807±0.074 | 0.844±0.130 |
|  |  | CT | 0.711±0.121 | 0.536±0.079 | 0.668±0.148 |
|  | *JSDs* | FD | 0.891±0.076 | 0.740±0.076 | **0.852±0.101** |
|  |  | GI | 0.766±0.109 | 0.605±0.075 | 0.728±0.140 |
|  |  | SD | **0.892±0.106** | **0.813±0.070** | 0.844±0.130 |
|  |  | CT | 0.709±0.121 | 0.537±0.078 | 0.668±0.148 |

The highest TRT reliability is highlighted in bold under each brain parcellation atlas and each analytical level. *KLDs*, Kullback-Leibler divergence-based similarity; *JSDs*, Jensen-Shannon divergence-based similarity; FD, fractal dimension; GI, gyrification index; SD, sulcal depth; CT, cortical thickness.

#### Effects of smoothing size

Based on the BNU TRT dataset, we found that the usage of a larger smoothing kernel (25-mm full width at half maximum) resulted in significantly higher TRT reliabilities for interregional morphological similarities and nodal centralities but lower TRT reliabilities for global network measures for the CT-based morphological similarity networks (paired t-test; *P* < 0.05, Bonferroni corrected; Table S3). Nonetheless, the differences did not affect our general findings that CT-based morphological similarity networks performed worse that the other three types of morphological similarity networks with respect to TRT reliability. That is, no matter which smoothing size (15-mm or 25-mm full width at half maximum) was used, significantly lower TRT reliabilities were found for the CT-based than FD-, GI- and SD-based morphological similarity networks for interregional morphological similarities, global network measures and nodal centralities (paired t-test; *P* < 0.05, Bonferroni corrected; Table S3). These findings indicate that differences in smoothing kernel among different morphological indices contribute little to the observed differences in TRT reliabilities among different types of morphological similarity networks.

### Relationships between morphological similarity networks and brain/regional size

Very low correlations were found between each global network measure and global morphological values regardless of the type of morphological similarity networks (Table S4). These findings suggest a lack of relationships between the morphological similarity networks and brain size. For nodal centrality measures, weak-moderate correlations were observed with regional size that were dependent on morphological index, brain parcellation atlas, similarity measure and nodal centrality type (Table S5). These findings indicate obvious effects of regional size on nodal centrality for the surface-based single-subject morphological similarity networks, an issue that should be taken into account in future studies. The findings of brain size-independent but regional size-dependent morphological similarity networks are in accordance to a previous study that constructed single-subject morphological similarity networks with different methods (Seidlitz et al., 2018).

## Discussion

In this study, we constructed and topologically characterized surface-based single-subject morphological similarity networks, and systematically evaluated their reproducibility with respect to effects of different analytical strategies, sample size-varying stability and test-retest reliability. We found that the morphological similarity networks exhibited nontrivial organizational principles, including small-worldness, high parallel efficiency, modularity and hubs, regardless of the analytical strategies used. Nevertheless, quantitative values of these organizational principles largely depended on the choices of morphological index, brain parcellation atlas and similarity measure. Moreover, the morphological similarity networks varied with the number of participants and approached stability until the sample size exceeded ∼70. Finally, both interregional similarities and topological properties of the morphological similarity networks presented fair to good, even excellent, TRT reliabilities. Interestingly, significantly higher reliabilities were observed when interregional morphological similarity was estimated with the *JSDs* than with the *KLDs* and when the morphological similarity networks were based on the FD and SD rather than the GI and CT. Altogether, these findings suggest that surface-based single-subject morphological similarity networks provide a reliable approach for large-scale brain network studies.

### Specifically organized and reliable morphological similarity networks

The human brain is powerful in both modular information processing across local regions and distributed information processing over the entire brain, allowing complicated cognition and behavior. A large number of studies have indicated that such powerful features of the human brain benefit from nontrivial wiring layouts of both functional and structural brain networks as revealed by graph-based network approaches, such as small-worldness, high parallel efficiency, modularity and hubs (Bullmore and Sporns, 2012; Liao et al., 2017; Sporns and Betzel, 2016; van den Heuvel and Sporns, 2013). Therefore, in this study we also utilized graph-based metrics to examine whether the surface-based single-subject morphological similarity networks constructed with our method are governed by nontrivial organizational principles rather than are wired in a random manner. We found that the surface-based single-subject morphological similarity networks also exhibited the optimized organization. That is, compared with matched random networks, the surface-based single-subject morphological similarity networks showed higher clustering coefficient, local efficiency and modularity, approximately equal characteristic path length and global efficiency and the existence of hubs. Moreover, the optimized organization was consistently observed regardless of different choices of morphological index, brain parcellation atlas and similarity measure. The robust and inherent characteristic indicates that the surface-based single-subject morphological similarity networks are specifically organized rather than are wired in a random manner. It should be noted that the interpretation of graph-based network findings largely depends on how network nodes and edges are defined. In this study, network edges were defined as divergence-based similarity in the distribution of intraregional morphological values. This type of similarity is very different from routinely used functional connectivity (i.e., statistical interdependence in functional signal fluctuations between regions) and structural connectivity (i.e., fiber pathways between regions). Thus, our findings are not directly comparable with previous functional and structural brain networks. It is interesting to investigate the relationships between the divergence-based morphological similarity and functional/structural connectivity in the future, which can aid in understanding the extent to which different modalities of brain networks are driven by similar organizational principles.

An interesting finding in this study was that consistent hub regions were mainly in cortical sulci under the a2009s atlas. Compared with cortical gyri, cortical sulci have been found to show greater physical strain in various kinds of injuries in studies from animals, computational models and human postmortem data (Cloots et al., 2008; Ghajari et al., 2017; Goldstein et al., 2012). Accordingly, cortical sulci are frequently reported to be more vulnerable to pathological tau protein deposition and associated brain atrophy in patients with traumatic brain injury and chronic traumatic encephalopathy (Cole et al., 2018; Johnson et al., 2012; McKee et al., 2013). On the other hand, brain network hubs are also found to be susceptible to various brain disorders (Crossley et al., 2014). Thus, our results provide a possible bridge to link the two sets of previous findings, although a direct examination is needed on the relationships among brain sulci and gyri, hubs and diseases. Finally, the surface-based single-subject morphological similarity networks presented fair to good, even excellent, reliability in both interregional similarities and network measures under all analytical combinations. This is consistent with previous studies that constructed individual-level morphological similarity networks with similar methods in brain volume space (Kong et al., 2015; Wang et al., 2016) and via different methods (Jiang et al., 2017; Li et al., 2017; Yu et al., 2018).

Taken together, our findings suggest that surface-based single-subject morphological similarity networks can provide an important alternative to reliably uncover organizational principles of large-scale brain networks. Nevertheless, it should be pointed out that although optimized topological organizations and high TRT reliabilities were consistently observed for the surface-based single-subject morphological similarity networks regardless of different analytical strategies, quantitative values depended largely on how the morphological similarity networks were constructed, which is discussed below.

### Effects of different morphological indices on morphological similarity networks

We found that different morphological indices resulted in significantly different values for interregional morphological similarities and network measures. This is consistent with previous studies of population-based morphological similarity networks showing morphological index specific topological organization in health and disease (Collantoni et al., 2017; Sanabria-Diaz et al., 2010). Given the highly complex, folded nature of the human cerebral cortex, these findings are not surprising because different indices characterize morphological architecture from different aspects. For example, CT reflects the size, density and arrangement of cells (neurons, neuroglia and nerve fibers), whereas GI measures the ratio between the total area of the cortex and the area that is visible in a circular region of interest. Thus, the observed differences may indicate that different morphological indices capture distinct processes of interregional interactions or different aspects of the same interactive process (e.g., mechanical, neurochemical, and/or axonal connections). Currently, understanding mechanisms behind different morphological indices and their interrelationships is an ongoing research field. Evidence from genetic and developmental studies has shown that different morphological indices are associated with distinct genetic influences (Panizzon et al., 2009; Strike et al., 2019; Winkler et al., 2010) and exhibit differential developmental/aging trajectories (Hogstrom et al., 2013; Raznahan et al., 2011; Wierenga et al., 2014). Accordingly, it is plausible to speculate that genetic, developmental/aging, and environmental factors may partly contribute to the distinct network topology among different morphological indices. It should be noted that the explanations above are speculative and more studies are needed to provide direct empirical evidence for mechanistic understanding of network differences among different morphological indices.

Furthermore, different morphological indices were associated with significantly different values of TRT reliability of the morphological similarity networks. Specifically, TRT reliability was significantly higher for FD- and SD- than for GI- and CT-based morphological similarity networks. The discrepancy may be due to differences in computational complexity, sensitivity to noise, and/or developmental rates among the morphological indices. For example, in contrast with the intuitional CT, FD is an extremely obscure and compact measure of shape complexity, which condenses all details into a single numeric value. Such summary measures may be more resistant to noise than those that are dominated by a single aspect of brain morphology. In addition, given distinct age-related trajectories among different morphological indices (Raznahan et al., 2011; Wierenga et al., 2014), those indices showing faster age-related changes are typically related to lower values of TRT reliability due to greater within-subject variance. Overall, our findings suggest that future studies should consider utilizing different morphological indices, in particular SD and FD from the perspective of TRT reliability, to provide a finer-grained characterization of surface-based single-subject morphological similarity networks in typical and atypical populations.

### Effects of different brain parcellation atlases on morphological similarity networks

We found that different choices of the a2009s and a2005s atlases resulted in significantly different surface-based single-subject morphological similarity networks in terms of the values of interregional morphological similarities and network measures. This is consistent with numerous previous studies showing brain parcellation-dependent functional (Ren et al., 2019; Wang et al., 2009), structural (Wei et al., 2017; Zalesky et al., 2010) and morphological (Sanabria-Diaz et al., 2010; Wang et al., 2016) brain networks (for recent reviews, see (Arslan et al., 2018; Qi et al., 2015; Yao et al., 2015)). Accordingly, our findings together with previous studies collectively suggest that the dependence on regional brain parcellation atlases is a universal characteristic of large-scale brain networks, and future studies must consider the influence of this factor regardless of the data modalities from which the brain networks are obtained. Several potential sources may account for the observed differences. First, the composition is different between the two atlases: the a2009s atlas is composed of sulco-gyral structures, while the a2005s atlas is made up of gyral-based neuroanatomical regions (Desikan et al., 2006; Destrieux et al., 2010). Given substantial differences between cerebral sulci and gyri in genetics, morphology, axonal pathways and function (Ge et al., 2018; Hilgetag and Barbas, 2005; Li et al., 2015; Liu et al., 2017; Zeng et al., 2015; Zhang et al., 2018), it is not surprising to observe differential network organization between the two atlases. Second, the number of regions is different between the two atlases: 148 in the a2009s atlas versus 68 in the a2005s atlas. Previous studies have shown that differences in network size (i.e., the number of nodes) significantly affect the topological organization of functional and structural brain networks (Wang et al., 2009; Zalesky et al., 2010). Finally, the two atlases differ in regional size, which is thought to be associated with nodal centralities of functional and structural brain networks (Hagmann et al., 2008; Wang et al., 2009). Together, all these factors may at least partially account for the observed brain parcellation-related differences in surface-based single-subject morphological similarity networks.

Further TRT reliability analyses revealed that the a2009s atlas was associated with higher reliability than the a2005s atlas for the surface-based single-subject morphological similarity networks. When using the high-resolution MMP atlas, even higher TRT reliability was observed. These findings indicate that brain parcellation atlases with higher resolutions may give rise to more reliable morphological similarity networks. The discrepancy may be attributable to the more sophisticated methods used to generate higher-resolution atlases: most structures in the a2005s atlas were defined using a relatively coarse ‘sulcal’ approach (manual tracing from the depth of one sulcus to another, thus incorporating the gyrus within) (Desikan et al., 2006); the entire cortical surface of the a2009s atlas was classified as gyral or sulcal at a vertex level during the generation (Destrieux et al., 2010); and the MMP atlas is based on a multi-modal method that integrates multidimensional information from different imaging modalities (e.g., architectural measures derived from T1-weighted and T2-weighted structural images, cortical function measured using task functional MRI, and functional connectivity estimated from resting-state functional MRI). Currently, obtaining accurate and reliable parcellation atlases of the human brain is an important goal for brain science. In this regard, individualized, multidimensional information guided and phylogeny and ontogeny inspired parcellation methods may be promising directions in the future (Eickhoff et al., 2018). Overall, our findings suggest that relative to the a2005s atlas and a2009s atlas, the MMP atlas may be a better choice for surface-based single-subject morphological similarity network studies from the perspective of TRT reliability.

### Effects of different similarity measures on morphological similarity networks

Although the morphological similarity networks constructed via *KLDs* and *JSDs* exhibited largely similar patterns, quantitative values differed significantly between the two sets of networks. This is consistent with numerous previous studies showing that functional and structural brain networks depend on the means to estimate interregional connectivity (Liang et al., 2012; Sarwar et al., 2019; Zalesky et al., 2012). Accordingly, it seems that the dependence on interregional connectivity estimation methods in addition to regional brain parcellation atlases is another universal characteristic of large-scale brain networks, and thus future studies should carefully choose suitable measures and methods for estimating interregional connectivity in terms of their research themes, purposes and contents. The observed differences may be attributable to different mathematical properties between the *KLD* and *JSD*, although the latter is based on the former. First, the *KLD* and *JSD* have different value ranges: the *KLD* is nonnegative without an upper limit, while the *JSD* is bounded by 0 and 1. Second, the *KLD* is not symmetric, while the *JSD* is symmetric. Thus, compared with the *JSD*, the *KLD* undergoes additional processing steps (e.g., exponential transform) to generate symmetric and bounded surface-based single-subject morphological similarity networks. This may introduce extra noise or unknown disturbance in estimating interregional morphological similarities. This might also be the reason why the *JSDs*-based morphological similarity networks showed higher reliability for interregional morphological similarities and network measures. Overall, given the mathematical advantages of the *JSD* relative to the *KLD* and higher TRT reliability of the *JSDs*-based than *KLDs*-based morphological similarity networks, we recommend using the *JSDs* as a measure for estimating interregional morphological similarity in future studies. In the future, it will be necessary to systematically compare the *JSDs* with other similarity measures and methods for a possible better choice, such as multivariate Euclidean distance (Yu et al., 2018) and Fréchet distance.

### Effects of different sample sizes on morphological similarity networks

We found that surface-based single-subject morphological similarity networks varied along with the number of participants and approached stability until the sample size exceeded ∼70. Moreover, the critical value was largely independent of different analytical strategies. Thus, we recommend at least 70 participants for future studies of surface-based single-subject morphological similarity networks. However, such a sample size may be challenging to attain for innovative clinical and translational research due to cost and feasibility concerns. Although small sample size undermines the reliability (Button et al., 2013), it can produce more projected scientific value per dollar spent than larger sample size for studies of new ideas (Bacchetti et al., 2011). Interestingly, we found that even when the sample size was only 10, the mean nodal centrality profile of the surface-based single-subject morphological similarity networks was largely similar to that from all 876 participants. This suggests limited effects of sample size on surface-based single-subject morphological similarity networks. This feature makes the approach proposed here a potential tradeoff between sample size and research cost and feasibility for future brain network studies. Presumably, the weak influence of sample size may reflect small interindividual differences in surface-based single-subject morphological similarity networks. An interesting topic for future exploration is the association of interindividual differences in surface-based single-subject morphological similarity networks with interindividual differences in cognition and behavior.

### Limitations and Future Directions

First, consistent with numerous functional and structural brain network studies, the surface-based single-subject morphological similarity networks were largely dependent on choices of regional brain parcellation atlases and similarity estimation methods. Thus, systematic research on different processing pipelines is warranted in the future for establishing a “better” methodological framework in the construction of surface-based single-subject morphological similarity networks. Second, although the surface-based single-subject morphological similarity networks exhibited high TRT reliability under different analytical strategies, their repeatability across multiple sites, scanners, scanning parameters and magnetic fields should be further examined. Third, similar to functional and structural brain networks, surface-based single-subject morphological similarity networks constructed here also exhibited optimized topological organizations (e.g., small-worldness and hubs). It will be interesting in the future to quantify the similarities and differences in organizational principles between morphological and functional/structural brain networks. Fourth, this study extended our previous work of single-subject morphological similarity networks from volume space to cerebral cortical surface. A previous functional MRI study showed that surface-based computation can increase TRT reliability of local short-range functional connectivity (Zuo et al., 2013). Whether surface-based morphological similarity networks outperform volume-based morphological similarity networks in TRT reliability is thus an interesting topic in the future. Fifth, only four surface-based morphological indices that were computationally available for the CAT12 toolbox were used to construct morphological similarity networks in this study. Future studies can examine the feasibility of our method on the basis of other morphological indices, such as surface area and surface normal. Sixth, in this study, we found that different morphological indices were associated with distinct interregional similarity patterns and topological organization of surface-based single-subject morphological similarity networks. It is important to develop network models to integrate the complementary information for a holistic view of morphological similarity networks. At the current stage, multilayer network methods may be a good solution for such a requirement (De Domenico, 2017; Vaiana and Muldoon, 2018). Seventh, we utilized binary rather than weighted network model to characterize morphological similarity networks in this study because weighted networks are computationally expensive. This is particularly important for this study because of the huge amount of computation. It’s essential for future studies to employ weighted network model, which can provide more information on the topological organization of morphological similarity networks, and may be more sensitive to capture morphological network alterations under conditions where interregional morphological similarities alter profoundly, such as diseases, development and aging. Finally, after demonstrating the reliable, nonrandom organization of the divergence-based single-subject morphological similarity networks, the next important thing is to uncover biological meaning of the networks. For example, to what extent are the morphological similarity networks under genetic control and to what extent do the morphological similarity networks determine individual behavioral and cognitive performance? Such studies are crucial to speed up future application of the divergence-based single-subject morphological similarity networks in health and disease.

## Conclusion

In conclusion, this study constructed and evaluated surface-based single-subject morphological similarity networks and demonstrated that the morphological similarity networks possessed nontrivial topological organization, were affected by different analytical strategies but largely independent of sample size, and exhibited high TRT reliability. Based on these findings, we conclude that the surface-based single-subject morphological similarity networks can serve as a reliable way to characterize large-scale brain networks in future studies.

## Data Availability Statement

All data that support the findings of this study are from publicly available datasets.

## Acknowledgements

This work was supported by the National Natural Science Foundation of China (Nos. 81922036 and 81671764), Key Realm R&D Program of Guangdong Province (No. 2019B030335001) and Key Realm R&D Program of Guangzhou (No. 202007030005). Data were provided [in part] by the Human Connectome Project, WU-Minn Consortium (Principal Investigators: David Van Essen and Kamil Ugurbil; 1U54MH091657) funded by the 16 NIH Institutes and Centers that support the NIH Blueprint for Neuroscience Research; and by the McDonnell Center for Systems Neuroscience at Washington University.

## Figure Legends

**Figure S1**. Hubs with the top 10% of nodal centrality of morphological similarity networks (brain parcellation: a2009s; similarity estimation: *JSDs*). FD, fractal dimension; GI, gyrification index; SD, sulcal depth; CT, cortical thickness; *k*_i_, nodal degree; *e*_i_, nodal efficiency; *b*_i_, nodal betweenness.

**Figure S2.** Mean similarity matrices and their intersession correlations and TRT reliability values for morphological similarity networks (brain parcellation: a2009s; similarity estimation: *JSDs*). No matter the choices of morphological index, the mean similarity matrices derived from session 1 (first row) and session 2 (second row) were highly correlated with each other (third row), with most connections showing good to excellent TRT reliability (i.e., *ICC* > 0.6, last row). *JSDs*, Jensen-Shannon divergence-based similarity; S1, session 1; S2, session 2; *ICC*, intraclass correlation; FD, fractal dimension; GI, gyrification index; SD, sulcal depth; CT, cortical thickness.

**Figure S3.** Connections showing significantly different values of TRT reliability between different morphological indices. Varied proportions of edges (numbers in white) showed morphological index-dependent TRT reliability, with a general pattern of FD > SD > GI > CT. *JSDs*, Jensen-Shannon divergence-based similarity; *KLDs*, Kullback-Leibler divergence-based similarity; FD, fractal dimension; GI, gyrification index; SD, sulcal depth; CT, cortical thickness.

**Figure S4**. Global organization of morphological similarity networks derived from the BNU TRT dataset (brain parcellation: a2009s; similarity estimation: *JSDs*). For both sessions, individual morphological similarity networks exhibited small-world, highly efficient and modular organization. FD, fractal dimension; GI, gyrification index; SD, sulcal depth; CT, cortical thickness.

**Figure S5**. TRT reliability values for global network metrics of morphological similarity networks (brain parcellation: a2009s; similarity estimation: *JSDs*). Most global metrics exhibited fair to excellent TRT reliability (i.e., *ICC* > 0.6) regardless of choices of morphological index. FD, fractal dimension; GI, gyrification index; SD, sulcal depth; CT, cortical thickness.

**Figure S6**. Global network measures showing significantly different TRT reliability values between different morphological indices. The TRT reliability of many global network measures depended on the choices of morphological index, with a general pattern of FD and SD > GI and CT. Squares/circles indicate significant/non-significant differences, and color bars indicate t-statistics. *JSDs*, Jensen-Shannon divergence-based similarity; *KLDs*, Kullback-Leibler divergence-based similarity; FD, fractal dimension; GI, gyrification index; SD, sulcal depth; CT, cortical thickness.

**Figure S7**. Spatial correlations of nodal TRT reliability maps under different analytical strategies. Under each parcellation atlas, high correlations were observed between nodal degree and efficiency and between different similarity measures; however, dramatically low correlations were observed between nodal betweenness and the other two nodal metrics and between different morphological indices. *ICC*, intraclass correlation; *JSDs*, Jensen-Shannon divergence-based similarity; *KLDs*, Kullback-Leibler divergence-based similarity; FD, fractal dimension; GI, gyrification index; SD, sulcal depth; CT, cortical thickness; *k*_i_, nodal degree; *e*_i_, nodal efficiency; *b*_i_, nodal betweenness; *, *P* < 0.05, Bonferroni corrected.

**Figure S8**. Regional maps at different levels of nodal TRT reliability (brain parcellation: a2009s; similarity estimation: *JSDs*). Most regions exhibited excellent TRT reliability for FD- and SD-based morphological similarity networks regardless of nodal measures. FD, fractal dimension; GI, gyrification index; SD, sulcal depth; CT, cortical thickness; *k*_i_, nodal degree; *e*_i_, nodal efficiency; *b*_i_, nodal betweenness.

**Figure S9**. Regions showing significantly different TRT reliability values between different morphological indices (brain parcellation: a2009s; similarity estimation: *JSDs*). Varied proportions of regions (numbers in white) were identified to show morphological index-dependent TRT reliability, with a general pattern of FD > SD > GI > CT. Similar patterns were also found under other combinations of brain parcellation atlas and similarity measure (Figure 18, 19 and 20). *JSDs*, Jensen-Shannon divergence-based similarity; FD, fractal dimension; GI, gyrification index; SD, sulcal depth; CT, cortical thickness; *k*_i_, nodal degree; *e*_i_, nodal efficiency; *b*_i_, nodal betweenness.

**Figure S10**. Regions showing significantly different TRT reliability values between different morphological indices (brain parcellation: a2009s; similarity estimation: *KLDs*). *KLDs*, Kullback-Leibler divergence-based similarity; FD, fractal dimension; GI, gyrification index; SD, sulcal depth; CT, cortical thickness; *k*_i_, nodal degree; *e*_i_, nodal efficiency; *b*_i_, nodal betweenness.

**Figure S11**. Regions showing significantly different TRT reliability values between different morphological indices (brain parcellation: a2005s; similarity estimation: *JSDs*). *JSDs*, Jensen-Shannon divergence-based similarity; FD, fractal dimension; GI, gyrification index; SD, sulcal depth; CT, cortical thickness; *k*_i_, nodal degree; *e*_i_, nodal efficiency; *b*_i_, nodal betweenness.

**Figure S12**. Regions showing significantly different TRT reliability values between different morphological indices (brain parcellation: a2005s; similarity estimation: *KLDs*). *KLDs*, Kullback-Leibler divergence-based similarity; FD, fractal dimension; GI, gyrification index; SD, sulcal depth; CT, cortical thickness; *k*_i_, nodal degree; *e*_i_, nodal efficiency; *b*_i_, nodal betweenness.

