## Supplemental Table S1 for "Surface-based Single-subject Morphological Brain Networks: Effects of Morphological Index, Brain Parcellation and Similarity Measure, Sample Size-varying Stability and Test-retest Reliability"

**Table S1.** Summary of the graph theoretical measures used in this study

| Measure | Character | Description |
| --- | --- | --- |
| **Global** | | |
| Clustering coefficient | *C*_p_ | The extent of local clustering or cliquishness of a network |
| Characteristic path length | *L*_p_ | The extent of overall routing efficiency of a network |
| Local efficiency | *E*_loc_ | The ability of parallel information propagation within local subgraphs or the extent of fault tolerance of a network |
| Global efficiency | *E*_glob_ | The ability of parallel information propagation through a network |
| Modularity | *Q* | The extent to which nodes in a network can be divided into subsets with dense edges within but sparse edges between them |
| **Nodal** | | |
| Degree | *k*_i_ | The number of edges linked to a node in a network |
| Efficiency | *e*_i_ | The efficiency by which a node communicates with other nodes in a network |
| Betweenness | *b*_i_ | The influence of a node over information flow among other nodes in a network |
