## Supplemental Table S2 for "Surface-based Single-subject Morphological Brain Networks: Effects of Morphological Index, Brain Parcellation and Similarity Measure, Sample Size-varying Stability and Test-retest Reliability"

**Table S2**. Spatial Pearson correlation coefficients in regional mean values between different morphological indices

|  | **a2009s atlas**  (148 nodes) | **a2005s atlas**  (68 nodes) |
| --- | --- | --- |
| FD vs GI | -0.253±0.080 | -0.271±0.102 |
| FD vs SD | 0.030±0.069 | 0.157±0.079 |
| GI vs SD | -0.299±0.085 | -0.346±0.085 |
| FD vs CT | -0.192±0.080 | -0.198±0.101 |
| GI vs CT | -0.476±0.083 | -0.501±0.083 |
| SD vs CT | 0.183±0.100 | 0.022±0.088 |

FD, fractal dimension; GI, gyrification index; SD, sulcal depth; CT, cortical thickness.
