## Supplemental Table S3 for "Surface-based Single-subject Morphological Brain Networks: Effects of Morphological Index, Brain Parcellation and Similarity Measure, Sample Size-varying Stability and Test-retest Reliability"

**Table S3.** Effects of smoothing size for CT maps on comparisons of TRT reliabilities between different types of morphological similarity networks

|  |  | Test-retest reliability | | |
| --- | --- | --- | --- | --- |
| Morphological index | Smoothing  size | Interregional similarity | Global  measure | Nodal  centrality |
| CT | 15 mm | 0.689±0.128 | 0.522±0.111 | 0.619±0.164 |
| CT | 25 mm | 0.715±0.120^a^ | 0.415±0.147^b^ | 0.649±0.162^a^ |
| FD | 25 mm | 0.889±0.066^a,c^ | 0.769±0.072^a,c^ | 0.838±0.114^a,c^ |
| GI | 25 mm | 0.753±0.116^a,c^ | 0.467±0.100^b,c^ | 0.691±0.162^a,c^ |
| SD | 25 mm | 0.861±0.123^a,c^ | 0.819±0.070^a,c^ | 0.839±0.142^a,c^ |

CT, cortical thickness; FD, fractal dimension; GI, gyrification index; SD, sulcal depth.

^a^Significantly higher than CT-based networks with 15-mm smoothing.

^b^Significantly lower than CT-based networks with 15-mm smoothing.

^c^Significantly higher than CT-based networks with 25-mm smoothing.
