## Supplemental Table S4 for "Surface-based Single-subject Morphological Brain Networks: Effects of Morphological Index, Brain Parcellation and Similarity Measure, Sample Size-varying Stability and Test-retest Reliability"

**Table S4.** Cross-subject Pearson correlation coefficients between global values of each morphological index and global network measures derived from corresponding morphological similarity networks

| Similarity  measure | Global  measure | a2005s/a2009s atlas | | | |
| --- | --- | --- | --- | --- | --- |
|  |  | FD | GI | SD | CT |
| JSDs-based networks | *C*_p_ | -0.002/-0.152 | 0.023/0.061 | 0.034/-0.098 | -0.019/0.027 |
|  | *L*_p_ | -0.034/-0.058 | 0.030/-0.106 | 0.005/0.081 | -0.053/0.037 |
|  | *E*_loc_ | 0.006/-0.145 | 0.020/0.061 | 0.021/-0.112 | -0.014/0.027 |
|  | *E*_glob_ | 0.037/0.059 | -0.031/0.098 | 0.009/-0.089 | 0.052/-0.037 |
|  | *Q* | -0.029/-0.106 | -0.001/0.000 | 0.058/-0.017 | -0.034/0.029 |
|  | nor *C*_p_ | -0.002/-0.152 | 0.023/0.061 | 0.034/-0.098 | -0.019/0.027 |
|  | nor *L*_p_ | -0.034/-0.058 | 0.030/-0.106 | 0.005/0.081 | -0.053/0.037 |
|  | nor *E*_loc_ | 0.006/-0.145 | 0.020/0.061 | 0.021/-0.112 | -0.014/0.027 |
|  | nor *E*_glob_ | 0.037/0.059 | -0.031/0.098 | 0.009/-0.089 | 0.052/-0.037 |
|  | nor *Q* | -0.029/-0.106 | -0.001/0.000 | 0.058/-0.017 | -0.034/0.029 |
| KLDs-based networks | *C*_p_ | -0.004/-0.153 | 0.026/0.061 | 0.026/0.061 | -0.022/0.028 |
|  | *L*_p_ | -0.033/-0.058 | 0.026/-0.110 | 0.026/-0.110 | -0.053/0.038 |
|  | *E*_loc_ | 0.003/-0.144 | 0.025/0.062 | 0.025/0.062 | -0.016/0.028 |
|  | *E*_glob_ | 0.037/0.059 | -0.027/0.101 | -0.027/0.101 | 0.052/-0.037 |
|  | *Q* | -0.034/-0.102 | 0.000/0.000 | 0.000/0.000 | -0.030/0.028 |
|  | nor *C*_p_ | -0.004/-0.153 | 0.026/0.061 | 0.026/0.061 | -0.022/0.028 |
|  | nor *L*_p_ | -0.033/-0.058 | 0.026/-0.110 | 0.026/-0.110 | -0.053/0.038 |
|  | nor *E*_loc_ | 0.003/-0.144 | 0.025/0.062 | 0.025/0.062 | -0.016/0.028 |
|  | nor *E*_glob_ | 0.037/0.059 | -0.027/0.101 | -0.027/0.101 | 0.052/-0.037 |
|  | nor *Q* | -0.034/-0.102 | 0.000/0.000 | 0.000/0.000 | -0.030/0.028 |

*JSDs*, Jensen-Shannon divergence-based similarity; *KLDs*, Kullback-Leibler divergence-based similarity; FD, fractal dimension; GI, gyrification index; SD, sulcal depth; CT, cortical thickness; *C*_p_, clustering coefficient; *L*_p_, characteristic path length; *E*_loc_, local efficiency; *E*_glob_, global efficiency; *Q*, modularity; nor, normalized.
