## Supplemental Table S5 for "Surface-based Single-subject Morphological Brain Networks: Effects of Morphological Index, Brain Parcellation and Similarity Measure, Sample Size-varying Stability and Test-retest Reliability"

**Table S5.** Person-level Pearson correlation coefficients between regional size and nodal centrality

| Similarity  measure | Morphological  index | Nodal  centrality | Nodal size | |
| --- | --- | --- | --- | --- |
|  |  |  | a2005s atlas | a2009s atlas |
| JSDs-based networks | FD | *k_i_* | 0.045±0.118 | -0.059±0.077 |
|  |  | *e_i_* | 0.079±0.119 | -0.016±0.073 |
|  |  | *b_i_* | -0.016±0.111 | -0.113±0.070 |
|  | GI | *k_i_* | 0.057±0.118 | -0.070±0.077 |
|  |  | *e_i_* | 0.116±0.106 | -0.035±0.078 |
|  |  | *b_i_* | -0.057±0.114 | -0.123±0.073 |
|  | SD | *k_i_* | 0.105±0.081 | -0.119±0.049 |
|  |  | *e_i_* | 0.129±0.074 | -0.049±0.047 |
|  |  | *b_i_* | -0.202±0.082 | -0.155±0.064 |
|  | CT | *k_i_* | 0.159±0.111 | -0.009±0.080 |
|  |  | *e_i_* | 0.163±0.099 | 0.019±0.079 |
|  |  | *b_i_* | 0.061±0.127 | -0.098±0.075 |
| KLDs-based networks | FD | *k_i_* | 0.043±0.117 | -0.062±0.077 |
|  |  | *e_i_* | 0.078±0.119 | -0.019±0.073 |
|  |  | *b_i_* | -0.016±0.111 | -0.113±0.070 |
|  | GI | *k_i_* | 0.054±0.118 | -0.072±0.077 |
|  |  | *e_i_* | 0.115±0.106 | -0.037±0.078 |
|  |  | *b_i_* | -0.058±0.114 | -0.122±0.073 |
|  | SD | *k_i_* | 0.103±0.081 | -0.121±0.049 |
|  |  | *e_i_* | 0.127±0.074 | -0.050±0.047 |
|  |  | *b_i_* | -0.201±0.081 | -0.157±0.064 |
|  | CT | *k_i_* | 0.159±0.111 | -0.012±0.080 |
|  |  | *e_i_* | 0.163±0.099 | 0.017±0.079 |
|  |  | *b_i_* | 0.061±0.127 | -0.098±0.075 |

*JSDs*, Jensen-Shannon divergence-based similarity; *KLDs*, Kullback-Leibler divergence-based similarity; FD, fractal dimension; GI, gyrification index; SD, sulcal depth; CT, cortical thickness; *k_i_*, nodal degree; *e_i_*, nodal efficiency; *b_i_*, nodal betweenness.
